# A structure-aware framework for genomic variant interpretation in genetic skeletal disorders

**DOI:** 10.64898/2026.03.15.711892

**Authors:** Serena G. Piticchio, Najla Hosseini, Giedre Grigelioniene, Laura Orellana

## Abstract

**Background:** Genetic skeletal disorders (GSDs) comprise a heterogeneous group of rare, predominantly monogenic conditions that are increasingly diagnosed through high-throughput sequencing. While gene discovery has progressed rapidly, interpretation of pathogenic and uncertain variants remains a major bottleneck, in part because their functional consequences are determined at the protein structure level.

However, a systematic assessment of structural knowledge across GSD-associated genes is currently lacking. Here, we present a comprehensive protein structure-centric analysis of 674 protein-coding genes implicated in GSDs.

**Methods:** We integrated experimental structures, AlphaFold2 (AF2) models, multimeric states, protein–protein interactions, and ClinVar variant annotations.

**Results:** We quantify experimental structural availability and sequence coverage, revealing that 37% of GSD proteins lack any experimental structure and that, among proteins with structures, sequence coverage is often incomplete. We show that AF2 models provide high-confidence structural information for a substantial subset of proteins lacking experimental data, but that model reliability strongly correlates with existing structural coverage. Analysis of multimeric assemblies and co-occurring partners demonstrates that many GSD proteins function as obligate multimers, highlighting the importance of interface-level interpretation of variants. Finally, mapping clinically annotated missense variants onto representative protein structures illustrates how structural context can inform the interpretation of pathogenic and uncertain variants, particularly at interaction interfaces.

**Conclusions:** Together, this work provides a structure-aware reference framework for GSD genes, highlighting systematic gaps in current protein knowledge and demonstrating how integration of structural data can guide genomic variant interpretation. Our observations support a broader principle of *structural equivalence*, whereby distinct variants converge on shared structural perturbations that explain clustering patterns and enable mechanistic interpretation of nearby variants of uncertain significance.

## Background

Genetic skeletal disorders (GSDs) [1,2] constitute a large and heterogeneous group of rare diseases characterized by abnormalities of bone and cartilage development, with clinical severity ranging from mild skeletal dysplasia to perinatal lethality. Although individual GSDs are often ultra-or hyper-rare [3], their combined incidence is estimated at approximately 1 in 2,000–5,000 births, making them among the most frequent categories of rare genetic disease. The recognition of the radiographic pattern phenotype is critically important in the diagnosis of GSD, and advances in massive parallel sequencing have dramatically improved molecular diagnostics. More than 500 genes have been implicated in GSDs to date. Nevertheless, a substantial fraction of diagnoses remains unsolved or only partially resolved at the level of variant interpretation.

A central challenge in clinical genomics is the interpretation of rare missense variants, many of which are classified as variants of uncertain significance (VUS) [4,5]. Current variant classification frameworks rely heavily on sequence-based features such as evolutionary conservation, population frequency, and predicted effects on protein stability or residue chemistry [6–9]. While these approaches are essential, they often fail to capture the deepest level at which amino acid substitutions exert their effects: the three-dimensional structure and conformational behaviour of proteins, which is key to understand biological function and central to evolutionary selection [10,11]. Because protein structure dictates molecular interactions, dynamics, and ultimately biological function, interpretation of genetic variation without structural context is inherently limited. Given the complexity of this task, only recently 3D-structures have started to be considered (see Missense 3D-DB, PhyreRisk, VarSite or AlphaMissense [12–15] for the simplest cases concerning mostly monomeric 3D-structures. However, proteins are typically part of large heterogeneous assemblies, which on one hand, makes mapping more challenging, and on the other, can reveal new insights on pathogenic mechanisms. For example, our structural mapping of variants in the ribosomal protein RPL13 revealed both pathogenic and VUS substitutions neatly clustered within a conserved RNA-binding Arg-fork motif located in one surface of a helix [40], rather than strict ribosomal assembly interfaces to other proteins. This pointed to disruption of extra-ribosomal regulatory functions involving mRNA binding in skeletal development as a core mechanism, beyond the previously assumed ribosomal dysfunction. Cases like RPL13 also emphasize how clustering onto shared motifs and interfaces can be “structurally equivalent” in terms of pathogenic outcome, as previously observed in the context of cancer [42,43]. Similar cross-mapping of cancer-associated variants and known Mendelian disease mutations can reveal unexpected structural overlaps that facilitate functional interpretation.[16].

Hence, protein structural data provide a powerful means to bridge the gap between genotype and phenotype by enabling spatial interpretation of variants within functional domains, active sites, and interaction interfaces. However, experimental structures deposited in the Protein Data Bank (PDB) are unevenly distributed across disease-associated genes and frequently represent only fragments of large or multidomain proteins. Moreover, many proteins function as obligate multimers or components of larger assemblies, yet variant interpretation is typically performed on monomeric structures, due to the complexity of structural and genomic database mapping and integration.

The advent of AlphaFold2 (AF2) [17,18] has substantially expanded the availability of predicted protein structures, offering near-complete proteome-wide coverage. While AF2 models provide unprecedented opportunities for structural interpretation of genetic variants [19,20], their reliability varies across proteins and regions, depending on factors such as sequence length, intrinsic disorder, and the availability of homologous structural templates. Systematic evaluation of where AF2 models can reliably complement experimental data, and where caution is warranted, is therefore essential before such models can be broadly integrated into clinical genomics workflows.

Our earlier work applying structural mapping pipelines to focused datasets provided multiple examples illustrating how structural context can inform variant interpretation [41]. In this study, we attempt the first systematic structural mapping of all variants in a disease group, extending our approach to protein assemblies and macromolecular complexes covering almost the complete known gene landscape of genetic skeletal disorders (GSDs). Our comprehensive, protein structure-centric framework for interpretation of mutations (**Figure 1**) integrates curated gene lists, experimental structures from the Protein Data Bank (PDB) [21,22], high-confidence AlphaFold2 models, multimeric assemblies, protein–protein co-occurrence data, and ClinVar variant annotations, with the aim to (i) quantify the current state of protein structural knowledge in GSD genes, (ii) identify systematic gaps in experimental and predicted structural coverage, and (iii) explore how structural context—particularly multimeric and interface-level information—can strengthen or even reframe the interpretation of rare variants in clinical genetics. Unlike previous studies focusing on individual disease genes or pan-cancer mutation datasets, this work systematically maps variants across an entire disease class, revealing recurring structural mechanisms shared among diverse molecular systems. To our knowledge, this represents the first systematic effort to integrate structural coverage, multimeric architecture, and clinical variant annotation across the full gene landscape of a defined Mendelian disease class.

**Fig. 1.**
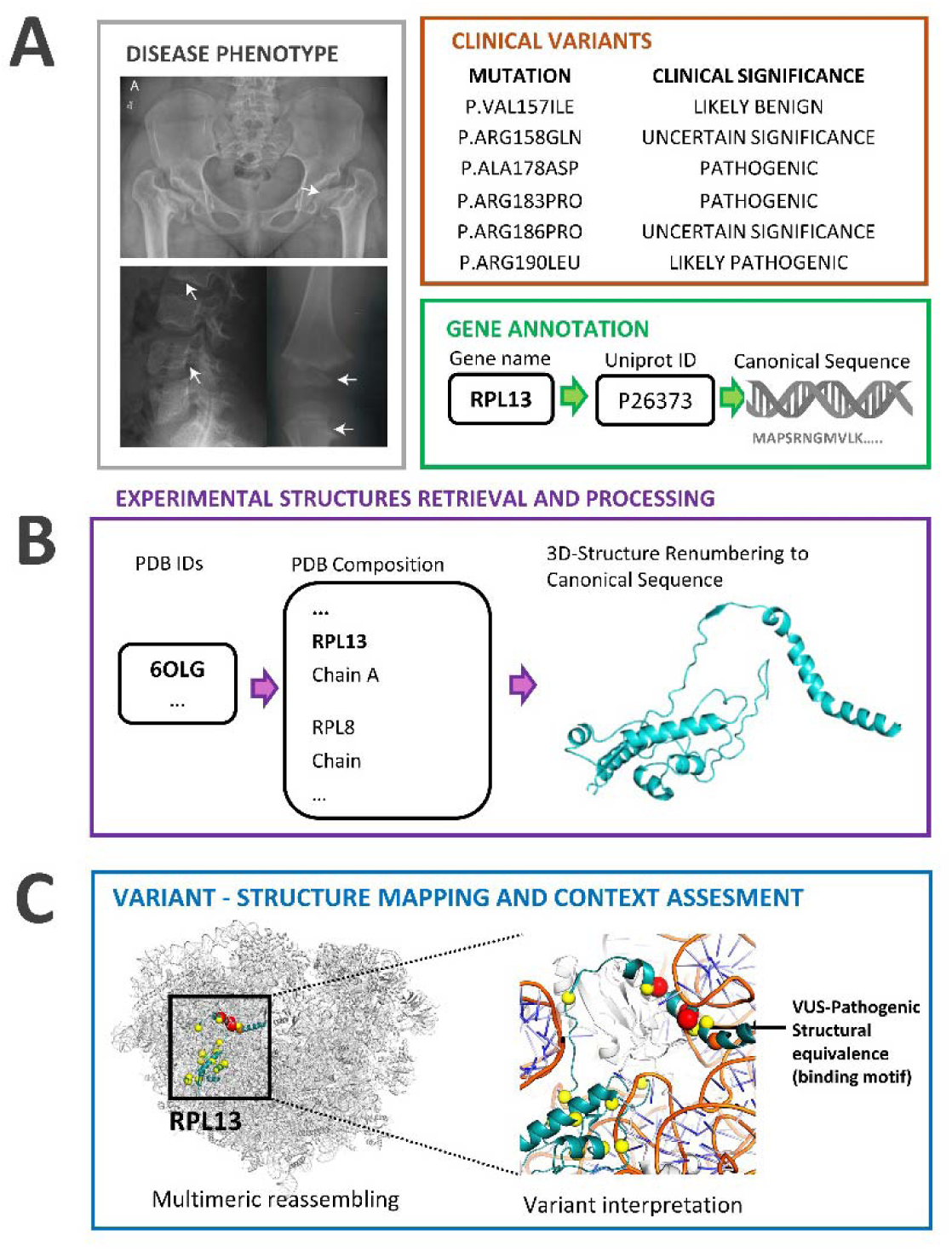
Structure-aware workflow for interpretation of disease variants and illustrative example from RPL13. Overview of the analytical workflow used in this study to integrate genomic and structural information for variant interpretation. Gene annotations (A) are first mapped to canonical UniProt sequences, followed by retrieval and processing of experimentally determined structures (B) from the Protein Data Bank (PDB). Protein chains are renumbered according to canonical sequences and reassembled to reconstruct multimeric complexes when necessary. Clinically annotated variants (e.g., from ClinVar) are then mapped onto representative protein structures (C) to evaluate spatial clustering within functional regions such as catalytic sites, RNA-binding motifs, or protein–protein interfaces. Structural mapping enables mechanistic interpretation of variants by revealing their proximity to interaction surfaces or functional motifs. As an illustrative example, missense variants in the ribosomal protein RPL13 cluster within a conserved RNA-binding helix that interacts with 28S rRNA in the human ribosome (PDB: 6OLG). Structural analysis previously demonstrated that these variants disrupt RNA binding rather than ribosome assembly per se, suggesting that the skeletal phenotype of SEMD-RPL13 may arise from perturbation of extra-ribosomal regulatory functions of eL13, including mRNA binding and translational control of pathways relevant to osteogenesis [40]. This example illustrates how structure-aware interpretation can generate mechanistic hypotheses that extend beyond conventional gene-centric disease classifications. Variants are colored by clinical significance: pathogenic (red), likely pathogenic (orange), variants of uncertain significance (yellow). In the example VUS (yellow) and a known pathogenic variant (red) map to the same positively charged RNA-binding helix surface.

## Methods

### Gene set definition and curation

A comprehensive list of genes associated with genetic skeletal disorders (GSDs) was assembled by integrating three independent datasets (**Supplementary Table S1**). The primary dataset was a curated GSD diagnostic gene panel provided by Karolinska University Hospital (hereafter “KI_PAN”), version 73.1 (November 2024), comprising 671 genes. This list was compared with the gene set reported in the 2023 revision of the Nosology of Genetic Skeletal Disorders [23] (“NOS23”). Genes with discrepant nomenclature were corrected **(Supplementary Table S2).** Genes present in KI_PAN but absent from NOS23 were retained. Genes listed in NOS23 but corresponding to loci, regulatory elements, or non-protein-coding entries were excluded (**Supplementary Tables S3**). Furthermore, a set of genes identified through clinical investigations of GSD patients at Karolinska University Hospital (“KI_CLI”) was added, including two genes present only in the clinical dataset (CDH11 and TOP3A) were added to the study. After consolidation and manual curation, the final dataset comprised 674 genes (**Supplementary Table S4 – Additional file 1**).

### Gene annotation and protein mapping

Gene symbols and HGNC identifiers [24] were retrieved from the source datasets when available. For genes lacking HGNC identifiers, manual annotation was performed using the HGNC database. Protein products were mapped to UniProt [25] reviewed human entries using the UniProt REST API [26], restricting queries to Homo sapiens (organism ID 9606) and reviewed entries. UniProt accessions were validated by cross-referencing HGNC identifiers and primary gene names. In cases of inconsistent nomenclature or alternative gene symbols, the UniProt primary gene name was adopted. Six genes in the curated dataset (*HAGLR, MIR140, MIR17HG, RMRP, RNU12, RNU4ATAC*) were identified as non-protein-coding and excluded from further analysis, as RNA structure and function were beyond the scope of this study (**Supplementary Table S5**). Two protein-coding genes (*GNAS, RAB34*) exhibited conflicting mappings between HGNC annotations, MANE Select transcripts [27], and UniProt canonical isoforms. Because these discrepancies prevented unambiguous reference sequence definition and residue numbering, these genes were excluded from structural analyses (**Supplementary Table S5**). Genes with ambiguous or conflicting annotations unrelated to skeletal disease (e.g. VCP, primarily associated with neurodegenerative disorders) were also excluded. (**Supplementary Table S5**). Additional protein annotations—including protein names and protein family classifications —were retrieved programmatically from UniProt using accession-based queries. The final protein-coding dataset comprised 674 genes (**Supplementary Table S6 – Additional file 1**).

### Reference proteome and canonical sequences

The human reference proteome (UniProt release October 2024; UP000005640_9606) was downloaded from the UniProt FTP repository (https://ftp.uniprot.org/pub/databases/uniprot/). Canonical protein sequences, as defined by UniProt, were used as reference sequences for residue numbering and sequence alignment. For each gene, the corresponding canonical FASTA sequence was extracted from the reference proteome using in-house scripts based on UniProt accession matching.

### AlphaFold2 structural models

Predicted protein structures were obtained from AlphaFold Protein Structure Database version 4 (AFDB-V4) [18, 28]. All human AF2 models were downloaded in November 2024. For proteins shorter than 2700 amino acids, a single full-length AF2 model was available. Proteins exceeding this length were represented by multiple overlapping models and were excluded from global AF2 confidence analyses due to ambiguity in reconstructing complete structures. Proteins lacking AF2 models were noted. For each AF2 structure, per-residue confidence scores (pLDDT) were extracted from the PDB files, and the average pLDDT score was calculated as a measure of overall model confidence. AF2 models were referenced using canonical UniProt residue numbering. Average confidence scores for all models are reported in **Supplementary Table S7 – Additional file 2**.

### Experimental structure retrieval, processing and sequence-coverage analysis

Experimentally determined protein structures were identified using the Structure Integration with Function, Taxonomy and Sequences (SIFTS) resource [29,30], which provides weekly updated mappings between UniProt accessions and PDB [21,22] entries. All associated PDB and mmCIF files were downloaded between January 31 and February 5, 2025, using in-house scripts based on the Biopython Bio.PDB module [31, 32]. Each structure was processed by separating individual protein chains into standalone files. For large complexes stored in mmCIF format, non-standard chain identifiers were normalized to ensure compatibility with PDB formatting. For each chain corresponding to a GSD protein, the observed amino acid sequence was extracted and aligned to the canonical reference sequence. Missing residues were annotated as unknown (“X”), and all structures were renumbered according to canonical UniProt residue numbering. The total number of experimental structures per protein is reported in **Supplementary Table S7 – Additional file 2**. Sequence coverage was defined as the fraction of residues in the canonical protein sequence that are present in at least one experimental structure. Coverage values were computed for all proteins with available experimental structures and used to stratify proteins into low-and high-coverage categories for downstream analyses.

### Pathway Analysis

Analysis was performed using Panther DB (https://pantherdb.org/) [33, 34] in December 2025. The Uniprot ID of all the 664 genes were pasted into the box and launched with “PANTHER Pathways” as the parameter in “Statistical overrepresentation test” analysis type and “Homo Sapiens” as species. “Homo Sapiens genes” was used as reference list. The parameters selected were “Fisher’s exact test” for test type and “False Discovery Rate (FDR)” for correction. 663 IDs were mapped to a Pathway, P02452 (COL1A1) was not mapped. All the results were downloaded and the pathways where at least one GSD gene is annotated are included in **Supplementary Table S10-S11 – Additional file 3**.

### Multimeric state and co-occurrence analysis

For each experimental structure, the number of chains corresponding to the protein of interest was recorded to assess observed oligomeric states. NMR ensembles were counted as a single structural instance. Proteins were classified based on whether they were observed exclusively as monomers, exclusively as multimers, or in both forms. To identify co-occurring proteins, all additional protein chains present in the same PDB entries were catalogued, irrespective of whether direct physical interactions were known. Instances where multiple GSD-associated proteins appeared within the same experimental structure were identified to reconstruct potential GSD-related complexes. Co-occurrence statistics and interaction groupings are reported in **Supplementary Table S12-S13 – Additional file 4**.

### Clinical variant retrieval and structural mapping

Clinically annotated variants were retrieved from the ClinVar database [5] in November 2024 only for assembly “GRCh38“. Variants were classified according to ClinVar clinical significance categories (**Supplementary Table S16)**. Missense variants were mapped onto representative experimental structures or AF2 models using canonical residue numbering. For illustrative structural analyses, a subset of proteins was selected based on high sequence coverage (≥95%), limited structural heterogeneity (≤3 experimental structures), and protein length (>400 residues) (**Supplementary Table S17)**. Variants were visualized in the context of monomeric or multimeric assemblies to assess spatial clustering and proximity to functional interfaces.

Cancer associated genes. The list of cancer associated genes, divided in Tier 1 and Tier 2 were downloaded from the Cancer Gene Census (CGC) webpage (https://cancer.sanger.ac.uk/cmc/home) [35] in July 2024 as text files with gene names in the first column. The gene names are compared with the gene names of the GSD list.

## Results

### Experimental structural knowledge biases and sequence coverage

The final curated dataset comprised 674 protein-coding genes associated with genetic skeletal disorders (GSDs) (**Supplementary Table S6 - Additional file 1 and Supplementary Figure 1**). Canonical protein lengths spanned more than two orders of magnitude, ranging from 55 amino acids (TOMM7) to 5,537 amino acids (KMT2D), with an average length of 803 residues (**Supplementary Figure 2**). This pronounced size heterogeneity reflects the diverse molecular roles of GSD proteins and foreshadows challenges in obtaining comprehensive experimental structural information, particularly for large, multidomain proteins. Experimental protein structures were available for 417 of the 664 proteins (63%), whereas 247 proteins (37%) lacked any experimentally determined structure (**Supplementary Figure 1 and Supplementary Figure 3**). Among proteins with structural data, representation was highly uneven. A small number of proteins—such as CA2, ESR1, and HRAS—were associated with hundreds of structures, reflecting intensive study in non-skeletal disease contexts, particularly cancer research.

Excluding these outliers, most proteins were represented by relatively few experimental structures, with an average of 18 structures per protein. It is worth noting that the presence of experimental structures did not imply comprehensive structural knowledge. Sequence coverage analysis revealed that only 105 of the 417 structured proteins (25%) had more than 90% of their canonical sequence represented, and just 19 proteins were fully covered (**Supplementary Figure 4**). In contrast, 150 proteins (35%) exhibited low coverage below 50%, including 32 proteins with coverage below 10%, often corresponding to isolated domains or short fragments captured within large complexes. The average coverage across all structured proteins was 62%, indicating that for most GSD genes, structural interpretation of variants is limited to specific regions rather than full-length architectures. Coverage deficits suggested differences across functional categories. Signalling enzymes and receptors often have higher coverage, whereas ECM proteins, transcriptional regulators, and large scaffolding proteins frequently displayed sparse or fragmentary structural representation. These gaps highlight systematic biases in experimental structural biology and have direct implications for variant interpretation.

### AlphaFold2 models in the context of experimental structures

AlphaFold2 (AF2) models were available for 648 of the 664 proteins. Proteins longer than 2,700 residues were excluded from global AF2 confidence analyses due to representation by multiple overlapping models, and one protein (GPX4) lacked an AF2 model. Across the remaining proteins, average AF2 confidence scores (mean pLDDT) ranged from 35.93 to 97.53, with an overall mean of 76.40 (**Supplementary Table S7**). When interpreted in isolation, AF2 confidence scores appeared broadly similar for proteins with and without experimental structures. However, stratification by experimental sequence coverage revealed a strong dependency of AF2 reliability on existing structural knowledge (**Supplementary Table S8**). Proteins with high experimental coverage (≥50%) exhibited consistently high AF2 confidence, with 93% showing average pLDDT scores ≥70. In contrast, proteins with low experimental coverage (<50%) exhibited markedly lower AF2 confidence: only 34% showed high-confidence models (mean pLDDT ≥70), with an average pLDDT of 65.62. Proteins entirely lacking experimental structures displayed an intermediate pattern, with 66% achieving high-confidence AF2 models. These findings indicate that AF2 models can provide valuable structural insight for many GSD proteins, including those without experimental data, but also underscore that prediction confidence is strongly influenced by underlying biological and structural constraints. In particular, proteins that depend on interaction partners or multimeric assemblies for stable folding often show reduced AF2 confidence when modelled as isolated monomers. The AF2 models evaluated here are predominantly monomeric predictions and therefore do not fully capture obligate multimeric interfaces or context-dependent conformational states. Many GSD-associated proteins function exclusively as components of higher-order assemblies, where folding, stability, or catalytic competence depends on inter-subunit interactions. In such cases, high per-residue confidence (pLDDT) in a monomeric model does not necessarily imply accurate representation of interface geometry or complex-dependent conformational rearrangements.

### Functional landscape of GSD-associated proteins

Functional annotation based on pathway enrichment analysis confirmed that GSD-associated proteins are heavily enriched in signaling and regulatory functions (**Supplementary Tables S10 - Additional File 3**) as previously observed. A substantial fraction participates in major developmental signaling pathways essential for skeletal formation and homeostasis, including Wnt, TGF-β/BMP, FGF, Hedgehog and Notch signaling. These signaling components are complemented by numerous transcription factors and transcriptional regulators, such as homeobox and zinc-finger proteins, which translate extracellular cues into gene expression programs during skeletogenesis. In addition, chromatin-associated and epigenetic regulators—including histone acetyltransferases, chromatin remodelers, and methylation-related proteins—are prominently represented, underscoring the importance of epigenetic control in skeletal development. Extracellular matrix (ECM) proteins, adhesion molecules, and enzymes involved in matrix modification form another major functional group, alongside transporters, metabolic enzymes, ribosomal proteins, and trafficking-associated factors. Together, these categories reinforce that GSDs arise from perturbations across multiple biological levels, from extracellular architecture to nuclear gene regulation.

### Structural interpretation of clinical variants and shared disease mechanisms

Analysis of experimental structures revealed that many GSD-associated proteins function within multimeric assemblies rather than as isolated monomers (**Supplementary Tables S12 and S13).** Among the 417 proteins with experimental structures, 82% were observed as monomers in at least one structure. However, 76 proteins were never observed as monomers and appeared exclusively in multimeric forms, indicating obligate oligomerization for functional stability or activity. Dimers were the most common multimeric state (61% of proteins with multimeric structures), followed by trimers (18%) and tetramers (32%). Larger assemblies were also observed, with some proteins captured in complexes containing more than 20 subunits, such as NLRP3 (21-mer) and CREBBP (24-mer). These observations underscore the importance of considering multimeric context when interpreting variants, particularly those located at protein–protein interfaces.

Co-occurrence analysis further demonstrated that most GSD proteins are captured experimentally alongside heterologous partners, reflecting participation in multi-protein complexes and signaling assemblies. Only 120 proteins lacked any hetero-partner across all structures, indicating that most GSD proteins are studied within multi-protein assemblies. Across 6,653 downloaded PDB entries, 442 contained more than one GSD-associated protein, corresponding to 115 unique GSD–GSD co-occurrence groups. These included 19 isolated protein pairs, 11 groups of three proteins, up to 13 larger interaction clusters, providing experimental support for shared molecular pathways underlying skeletal phenotypes.

Mapping ClinVar missense variants onto representative protein structures illustrated the functional relevance of these interaction contexts (**Figures 2 and 3**). Representative proteins were selected based on near-complete structural coverage (≥95%), limited conformational heterogeneity, and sufficient length to capture complex architectures, allowing reliable visualization of variant distributions within structurally defined regions (Supplementary Table S17). Mapping of pathogenic and likely pathogenic variants often revealed clustering within functional domains and, notably, suggested that interface-proximal mutations represent a recurrent mechanism in GSDs. In several proteins, variants classified as VUS localized in close proximity to known pathogenic substitutions within structurally defined functional regions. Such spatial proximity suggests that some VUS may represent under-classified pathogenic variants, particularly when the associated clinical phenotype is strongly consistent with known genotype–phenotype correlations. As mentioned above, a clear example is SEMD-RPL13, where pathogenic variants and VUS appear neatly clustered on Arg-fork on the surface of a helix and thus are essentially “equivalent” in structural terms (**Figure 1**). These substitutions disrupt a RNA recognition motif broadly found beyond strict ribosome assembly, suggesting that the skeletal phenotype may arise from perturbation of extra-ribosomal regulatory functions of RPL13 [40], including binding to specific mRNA of transcription factors involved in skeletal development (NF-κB). Another clear example is provided by BBS1, a core component of the BBSome complex. Structural mapping shows that a pathogenic BBS1 variant localizes in close proximity to the interaction interface with BBS4 within the assembled BBSome (**Figure 4**).

**Fig. 2.**
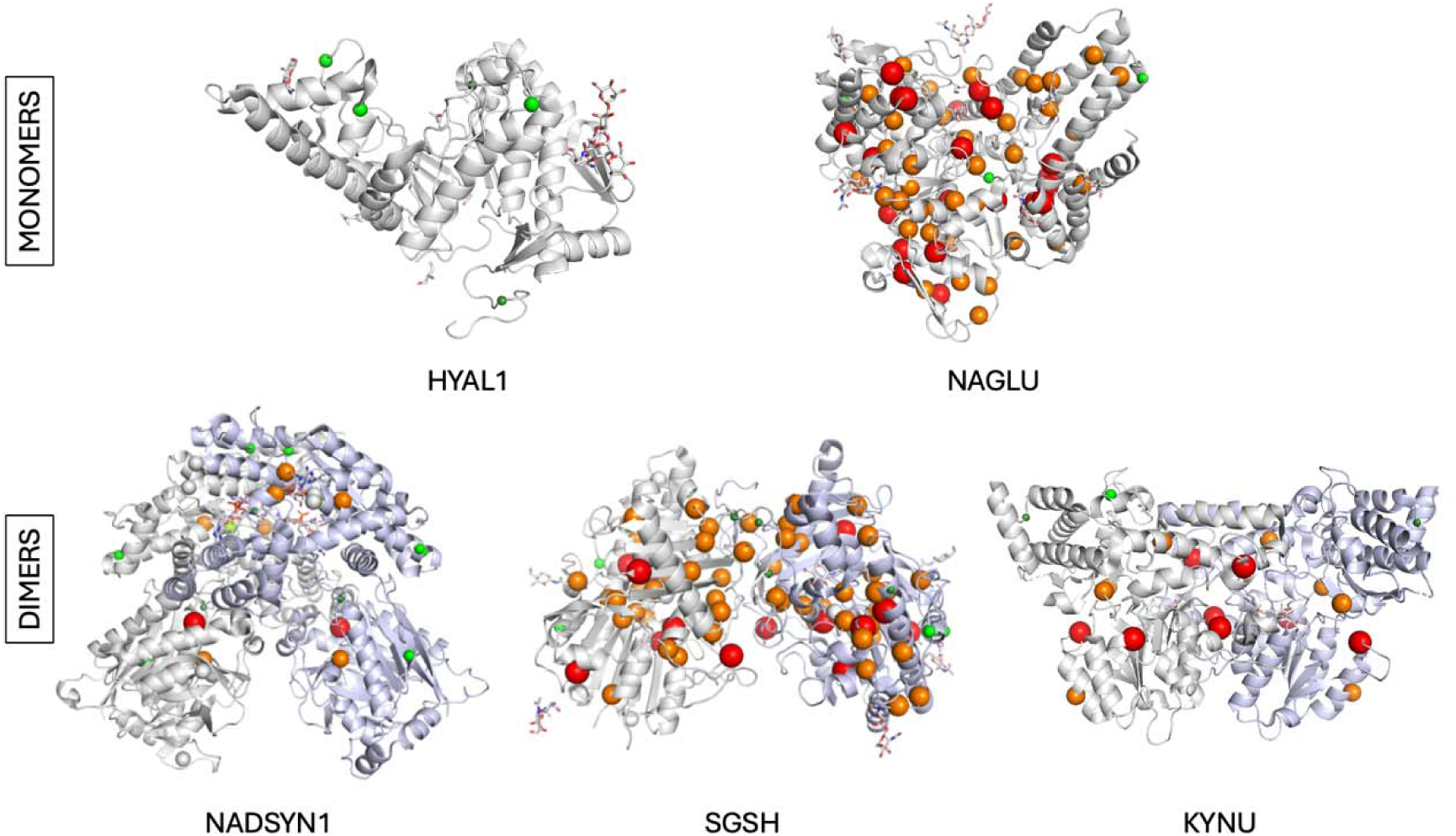
Representative structural mapping of ClinVar missense variants in monomeric and homodimeric proteins reveals clustering in functional domains and near pathogenic substitutions. Representative monomeric or homodimeric proteins with ClinVar non-VUS missense variants mapped onto experimental structures: HYAL1 (PDB code 2pe4), NAGLU (PDB code 4xwh), NADSYN (PDB code 6ofb), SGSH (PDB code 4mhx), KYNU (PDB code 2hzp). Variants are colored by clinical significance: pathogenic (red), likely pathogenic (orange), likely benign (light green), and benign (green). Mapping reveals clustering of pathogenic and likely pathogenic variants within functional domains and highlights regions sensitive to substitutions.

**Fig. 3.**
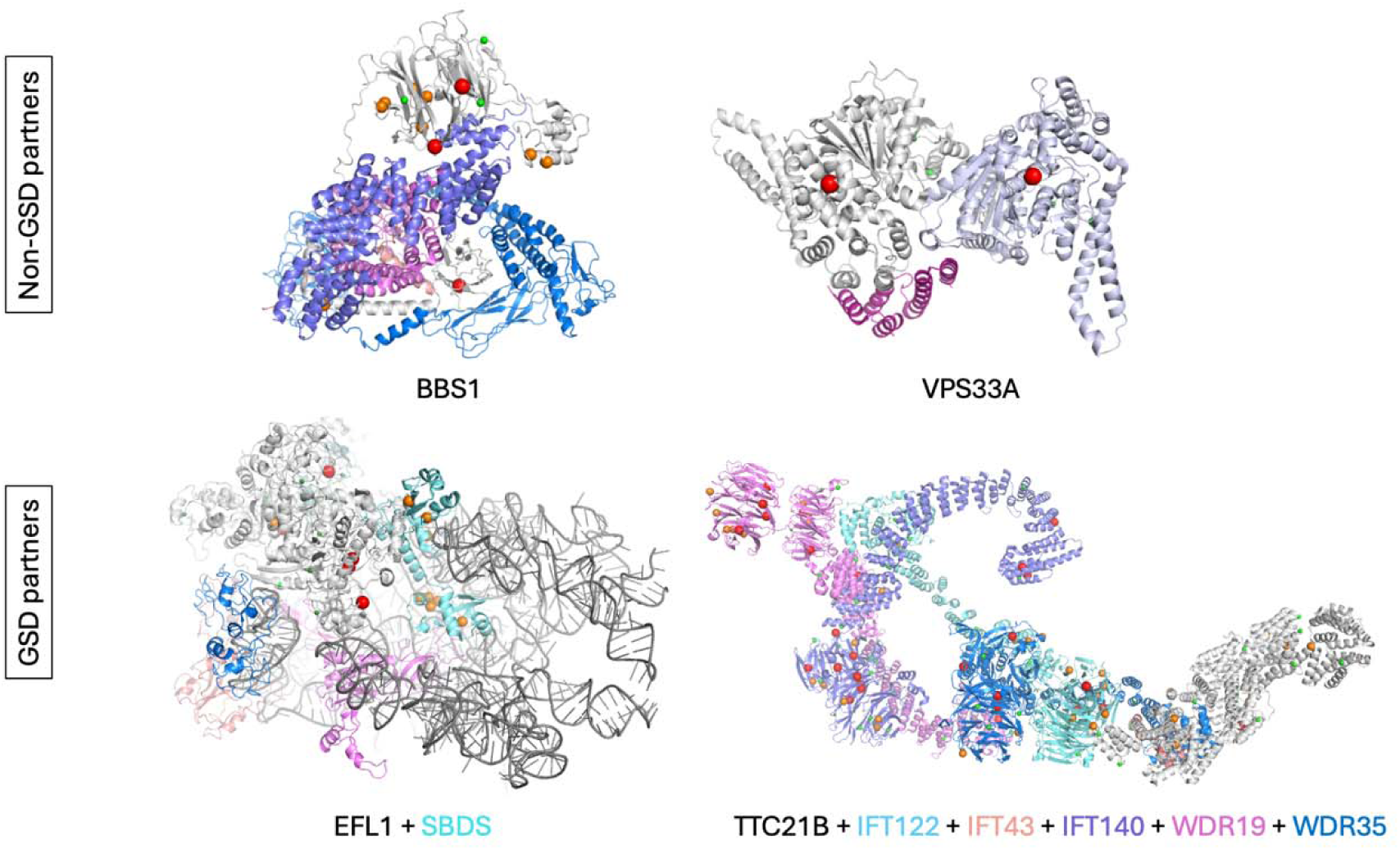
Structural mapping of ClinVar missense variants onto heteromeric protein assemblies reveals clustering at interaction interfaces. ClinVar missense variants mapped onto heteromeric protein complexes involving GSD-associated proteins and interaction partners not associated currently with GSD: BBS1 (PDB code 6xt9), VPS33A (PDB code 4bx9), EFL1 (PDB code 5anb), TTC21B (PDB code 8bbg). Structural context shows enrichment of pathogenic and likely pathogenic variants at protein–protein interacting interfaces, emphasizing the importance of complex-level interpretation for accurate variant classification. Colors according to pathogenicity (Red: Pathogenic; Orange: Likely Pathogenic; Light Green: Likely Benign; Green: Benign).

**Fig. 4.**
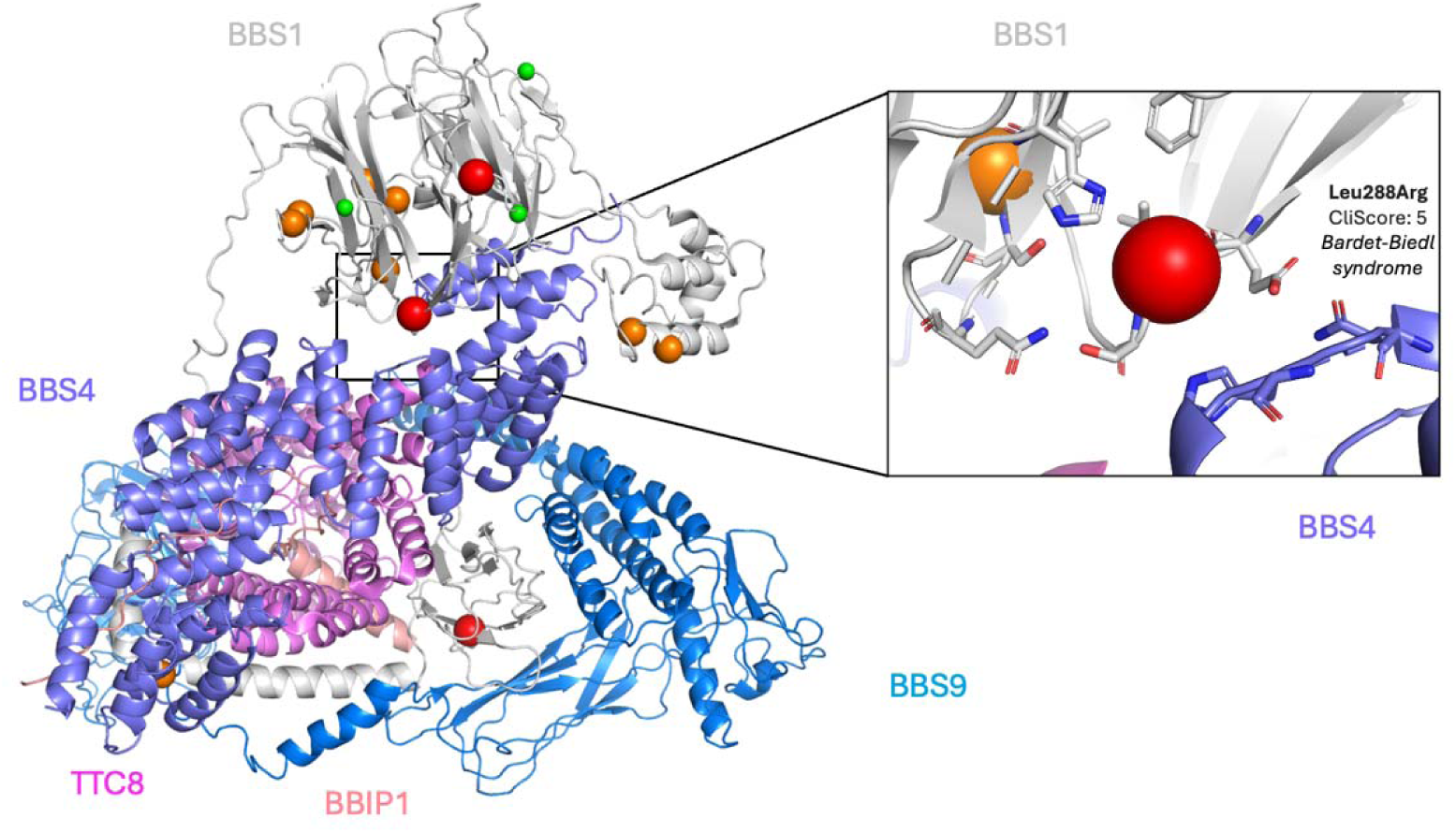
Interface-localized pathogenic variant in the BBSome complex potentially disrupting assembly. Structural representation of BBS1 within the BBSome complex (PDB ID: 6XT9), highlighting a pathogenic BBS1 variant located adjacent to the interaction interface with BBS4. Disruption of this interface provides a mechanistic explanation for impaired BBSome assembly and function, illustrating how multimer-aware structural mapping informs pathogenicity. Variants are colored by clinical significance: pathogenic (red), likely pathogenic (orange), likely benign (light green), and benign (green).

Given the obligate nature of this interaction for BBSome stability and trafficking function, disruption of the BBS1–BBS4 interface provides a direct structural explanation for pathogenicity through impaired complex assembly rather than isolated loss of BBS1 folding or stability.

Pre-existing experimental validation of interface sensitivity is also illustrated by VPS33A, a component of the HOPS tethering complex [36]. Several ClinVar variants of uncertain significance (VUS) cluster near the VPS33A–VPS16 interaction surface. Notably, two residues at this interface, Tyr438 and Ile441, have been experimentally shown through mutagenesis to be critical for VPS16 binding, with substitutions at these positions disrupting complex formation in vitro [37]. Structural mapping thus provides retrospective validation that interface-localized VUSes can be functionally deleterious even when sequence-based predictors are inconclusive (**Figure 5**). A further illustration is provided by the MCM helicase complex, which includes eleven proteins associated with genetic skeletal disorders, particularly Meier–Gorlin syndrome. Because MCM proteins function exclusively as a obligate heteromeric assembly, variants must be interpreted in the context of the full complex rather than individual subunits. Structural mapping across the assembled complex reveals that the known pathogenic Meier–Gorlin syndrome variant Thr466Ile in MCM5 localizes to the catalytic site formed at the interface with MCM3. Within the same catalytic region of MCM3, multiple ClinVar-classified VUSes (Arg356 and Thr370) are present. Given their shared spatial localization at the inter-subunit catalytic interface, it is likely that these MCM3 variants impair similarly helicase activity by disruption of ATP hydrolysis (another potential instance of structural-functional equivalence), and may represent pathogenic alleles that are currently under-classified (**Figure 6**). Together, these examples demonstrate that interface-aware structural mapping enables mechanistic interpretation of both pathogenic variants and VUSes, particularly in proteins that function as obligate multimers. Such analyses reveal pathogenic mechanisms—such as disruption of complex assembly or inter-subunit catalysis—that are not captured by monomer-centric or even less sequence-only approaches, underscoring the importance of incorporating quaternary structure and motif annotations into clinical variant interpretation. While the examples presented here illustrate interface-localized pathogenic variants, the present study was designed to establish a structural landscape rather than perform exhaustive mutational enrichment analyses. Future work integrating quantitative interface annotations with large-scale variant datasets will be required to evaluate interface enrichment across the full GSD protein set.

**Fig. 5.**
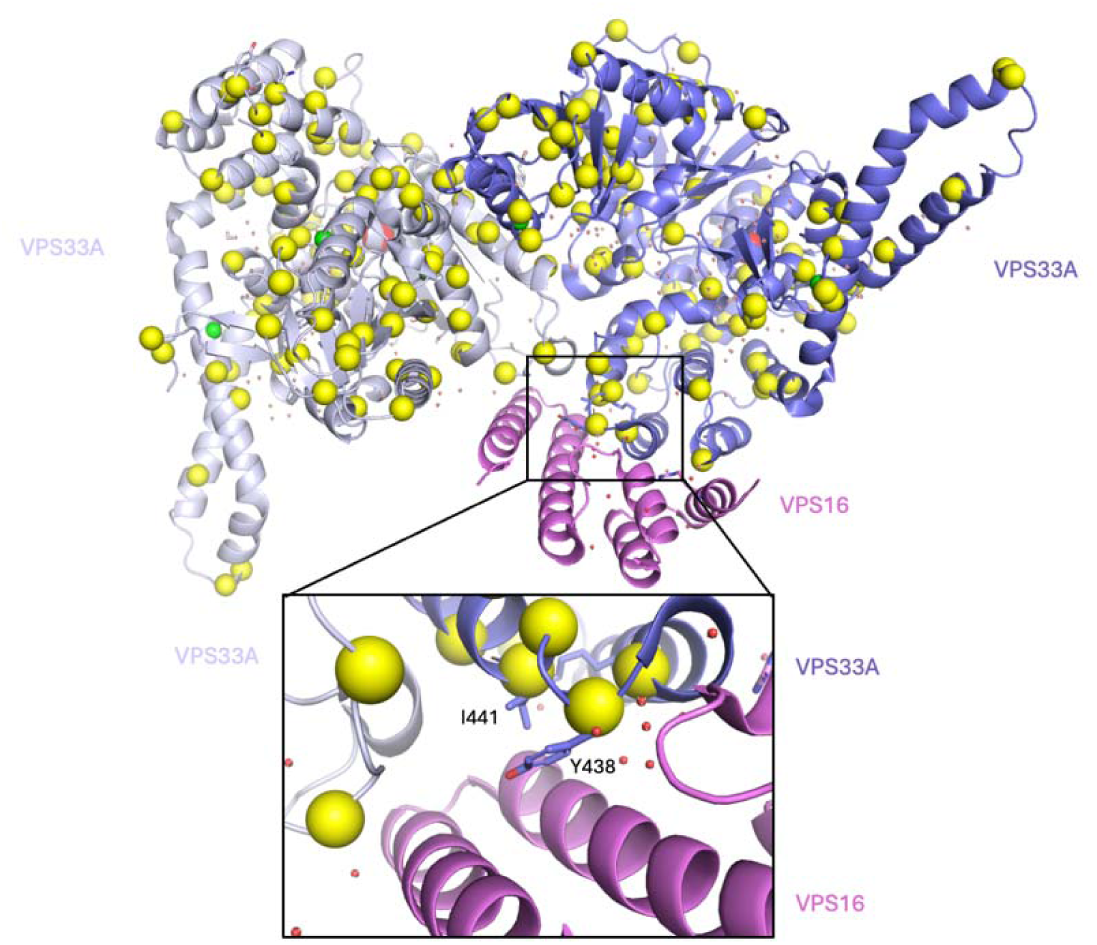
Structural context of VPS33A variants at the VPS33A–VPS16 interface within the HOPS complex disrupting assembly. Crystal structure of VPS33A in complex with its interaction partner VPS16, a core component of the HOPS tethering complex (PDB ID: 4BX9). VPS33A is shown in cartoon representation, with interface residues highlighted. ClinVar missense variants mapped onto VPS33A are colored by clinical significance: pathogenic (red), likely pathogenic (orange), variants of uncertain significance (yellow), likely benign (light green), and benign (green). Two interface residues, Tyr438 and Ile441, previously shown by mutagenesis to disrupt VPS16 binding when substituted, localize within the same interaction surface. This structural context supports the interpretation that interface-proximal variants can impair complex assembly and underlie pathogenic mechanisms in genetic skeletal disorders.

**Fig. 6.**
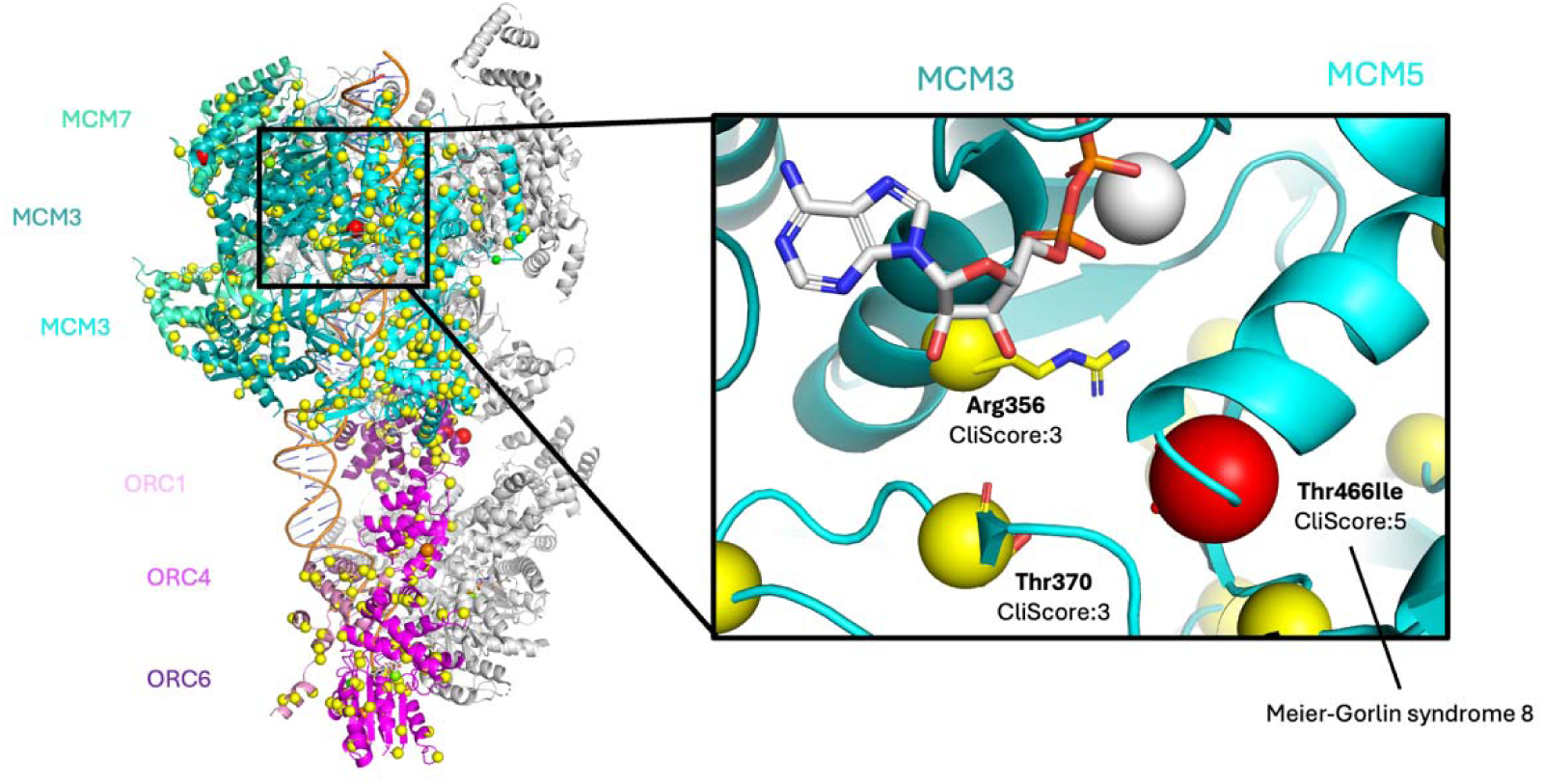
Structural mapping of ClinVar missense variants across the MCM helicase complex, formed by multiple proteins implicated in genetic skeletal disorders. The known pathogenic Meier–Gorlin syndrome variant Thr466Ile in MCM5 localizes to the catalytic site formed at the interface with MCM3. Variants of uncertain significance in MCM3 cluster within the same inter-subunit catalytic region, suggesting potential pathogenic effects that are not evident from sequence-based analysis alone. Variants are colored by clinical significance according to ClinVar pathogenic score (Red: Pathogenic; Orange: Likely Pathogenic, Yellow: Uncertain Significance; Light Green: Likely Benign; Green: Benign)

### Overlap with cancer-associated genes and implications for variant interpretation

Interestingly, among GSD genes are well-studied cancer genes such as FGFR1-3, CREBPP, MAP3K7, KRAS, NOTCH2, IDH1, PIK3R1 and others. Cross-referencing the GSD protein dataset with a publicly available snapshot of the Cancer Gene Census identified up to 102 genes shared between GSDs and cancer-associated gene sets (**Supplementary Table S18**). These overlapping genes were strongly enriched in signalling pathways, transcriptional regulation, and chromatin modification, including components of the PI3K–AKT, RAS/MAPK, Notch, and receptor tyrosine kinase pathways, as well as epigenetic regulators such as CREBBP, EP300, ARID1A, and SMARCA4. This overlap reflects however shared molecular principles rather than clinical phenotypes, although examples of potential functional overlap between cancer mutations and those of rare diseases have been observed [41]. Genes that govern cell fate decisions, proliferation, and transcriptional plasticity are particularly sensitive to functional perturbation: somatic mutations in these genes can drive oncogenesis, whereas germline variants disrupt tightly regulated developmental programs, leading to skeletal disorders. Importantly, cancer genomics has long leveraged structural clustering and interface-level interpretation of variants [38,39]. The presence of these shared genes underscores the potential to transfer structural insights and interpretative strategies from cancer research to the analysis of rare germline variants in GSDs.

## Discussion

High-throughput sequencing has transformed the diagnosis of genetic skeletal disorders (GSDs), yet interpretation of rare and novel variants remains a major unresolved challenge. While extensive gene catalogues and variant databases exist, the functional consequences of genetic variation are ultimately determined at the protein level, through effects on structure, interactions, and molecular mechanisms. In this study, we present a comprehensive, protein-centric landscape of GSD-associated genes that integrates experimental structural knowledge, AlphaFold2 (AF2) models, multimeric assemblies, biological function, and clinical variant data. By systematically linking genomic variation to protein structure and context, our work provides a framework for improving variant interpretation in skeletal disorders and, more broadly, in rare disease genomics.

A central finding of this study is that structural knowledge across GSD proteins is highly uneven and systematically biased. More than one-third of GSD-associated proteins lack any experimentally determined structure, and even among proteins with structural data, sequence coverage is frequently fragmentary. These gaps are not randomly distributed: large multidomain proteins, extracellular matrix components, transcriptional regulators, and scaffolding proteins—despite their central roles in skeletal development—are disproportionately underrepresented in structural databases. In contrast, signaling enzymes and receptors, particularly those studied extensively in oncology and drug discovery, tend to have higher structural coverage. This imbalance has direct implications for genomic interpretation, as absence of structural information limits the ability to assess the functional impact of missense variants, especially in ultra-rare conditions where statistical or cohort-based evidence is unavailable.

The availability of AlphaFold2 models offers an unprecedented opportunity to extend structural insight across the proteome, and our analysis demonstrates that AF2 can indeed provide high-confidence structural information for a substantial subset of GSD proteins lacking experimental data. However, our results also highlight important limitations. AF2 confidence strongly correlates with existing experimental coverage, thus carrying the highlighted biases in experimental structural data. In general, proteins with low structural coverage or obligate dependence on interaction partners frequently exhibit reduced model confidence. This suggests that AF2 predictions are most reliable when embedded within an informed biological context and should not be interpreted in isolation. Recent advances in structure prediction are beginning to address some of these limitations. In particular, AlphaFold-Multimer and the recently introduced AlphaFold3 framework extend prediction capabilities to protein–protein interactions and larger macromolecular assemblies. These developments hold promise for improving structural interpretation of variants located at interaction interfaces or within multi-protein complexes. However, such approaches remain computationally demanding and are not yet routinely integrated into large-scale clinical genomics pipelines. For the foreseeable future, reliable interpretation of interface-localized variants will therefore continue to rely on careful integration of predicted models with experimentally resolved assemblies and biological context. Overall, this underscores the need for cautious, context-aware evaluation of predicted structures rather than blanket reliance on confidence scores alone.

Consistent with these emerging structural prediction developments, a key insight emerging from our analysis is the central role of multimeric assemblies and interaction interfaces in the molecular pathology of GSDs. Many GSD-associated proteins function as obligate oligomers or as components of larger complexes, and a significant fraction are never observed experimentally as monomers. Mapping clinically annotated missense variants onto representative structures reveals that pathogenic variants often localize to protein–protein interfaces or shared catalytic surfaces formed by multiple subunits. Such variants may exert dominant-negative or complex-disrupting effects that are difficult to predict using sequence-based approaches alone. These observations argue strongly for moving beyond monomer-centric models of variant interpretation and incorporating complex-level structural information into diagnostic workflows. Although emerging approaches, including AlphaFold-Multimer and next-generation structure prediction frameworks, aim to address protein–protein complex modelling, systematic integration of multimeric predictions into clinical variant interpretation remains limited and still challenging. These considerations reinforce the need to interpret predicted models within known quaternary and pathway contexts, particularly when assessing variants located near obligate interaction surfaces.

The overlap between GSD-associated genes and cancer-associated genes further reinforces this perspective. A substantial subset of GSD genes also appears in cancer gene catalogues, particularly among regulators of signal transduction, transcription, and chromatin organization. Although the clinical manifestations of germline skeletal disorders and somatic cancers are distinct, the underlying molecular logic is shared: genes that govern cell fate decisions, proliferation, and transcriptional plasticity are highly sensitive to functional perturbation. In cancer genomics, structural clustering and interface-level interpretation of variants have long been used to identify driver mutations and infer mechanisms. Our findings suggest that similar structure-aware strategies could be fruitfully applied to rare germline disorders, enabling cross-disciplinary transfer of interpretative approaches.

From a translational perspective, the framework presented here has several implications for clinical genomics. Structure-aware annotation can help prioritize variants of uncertain significance by revealing proximity to functional domains, interaction interfaces, or conserved structural features. It can also suggest plausible mechanisms—such as loss of function, dominant-negative interference, or altered complex assembly—that guide downstream functional assays. Importantly, structural interpretation can also challenge existing disease classifications. As mentioned above, analysis of SEMD-RPL13 variants revealed clustering in an RNA-binding motif of the ribosomal protein eL13 [40], suggesting broader disruption of extra-ribosomal regulatory functions. Moreover, RPL13 illustrates how different amino acid substitutions clustered at the same structural motif can produce a shared functional consequence. A similar principle of structural equivalence has been described in oncogenic mutations of EGFR in glioblastoma [42,43], where structurally heterogeneous alterations converge on a shared conformational intermediate by removing a common steric constraint, illustrating how distinct mutations can produce equivalent functional outcomes. Recognition of such structurally equivalent perturbations provides a practical framework for variant interpretation, as variants of uncertain significance that cluster within the same structural motif or interface as known pathogenic substitutions may reasonably be inferred to affect the same molecular mechanism. In SEMD-RPL13, pathogenic variants cluster within the RNA-binding surface of helix H7, where conserved arginine residues participate in an “arginine-fork” motif that stabilizes RNA hairpin structures through interactions with the phosphate backbone. Mutations affecting these residues therefore represent structurally equivalent perturbations that converge on disruption of RNA binding, supporting a mechanism involving altered RNA recognition rather than strict ribosomal dysfunction.

Our earlier work applying a preliminary version of our structural mapping in a cohort of fetal skeletal dysplasias analyzed by genome sequencing, allowed the interpretation of multiple pathogenic variants through their impact on protein structure and functional domains. For instance, substitutions in collagen genes such as COL1A1, COL1A2, and COL2A1, which encode core components of the extracellular matrix, typically occur within the glycine-rich repeats that form the collagen triple helix. Replacement of glycine residues within the Gly-X-Y motif disrupts the tight packing required for triple-helix formation, leading to destabilization of collagen fibrils and impaired cartilage matrix organization. Similarly, variants in enzymes such as ALPL, encoding tissue-nonspecific alkaline phosphatase, affect residues involved in structural stabilization of catalytic loops and dimerization interfaces, thereby impairing enzymatic activity required for bone mineralization. These examples showed how structural interpretation can provide mechanistic support for variant classification even in the absence of functional assays.

The current framework extends this principle by systematically mapping variants onto protein assemblies and macromolecular complexes involved in diverse cellular pathways. Several examples highlighted in our figures illustrate how pathogenic variants affect proteins operating within higher-order functional modules. Components of the HOPS complex, for example, participate in vesicle tethering and fusion events between late endosomes, autophagosomes and lysosomes, processes that are critical for intracellular trafficking and turnover of cellular components. Variants affecting proteins within this pathway may impair lysosomal degradation and secretion dynamics that are essential for cartilage matrix maintenance. Likewise, variants affecting components of the BBSome complex implicate defects in ciliary trafficking, a process required for proper localization of signaling receptors within the primary cilium and for the regulation of developmental pathways such as Hedgehog signaling that are central to skeletal patterning. Finally, variants affecting proteins within the MCM helicase complex, which forms the core replicative helicase responsible for DNA unwinding during replication, highlight the importance of genome replication and proliferative capacity in rapidly dividing growth-plate chondrocytes. Together, these examples illustrate how structural mapping can place pathogenic variants within the functional architecture of cellular systems—ranging from extracellular matrix assembly and metabolic enzymes to vesicle trafficking, ciliary signaling, and DNA replication—providing a mechanistic framework for interpreting the diverse genetic causes of skeletal disease. Notably, some interacting partners within these complexes have not yet been linked to GSD, but their structural and functional dependence on the same assemblies suggests that variants affecting these proteins could produce related skeletal phenotypes.

Importantly, this approach is particularly valuable for ultra-rare and hyper-rare conditions beyond GSDs, where traditional criteria for pathogenicity based on recurrence or segregation are often unattainable. By integrating structural context into genomic interpretation, it becomes possible to extract mechanistic insight even from single-patient observations.

The integration of protein structural context into clinical genomics requires not only conceptual relevance but practical alignment with established variant classification frameworks. In current clinical practice, variant interpretation is largely guided by ACMG/AMP criteria, which emphasize population frequency, segregation, computational predictions, and functional evidence. However, structural context is often incorporated in a fragmented or informal manner, typically limited to assessing whether a variant lies within a known functional domain. Our systematic analysis suggests that a more thorough incorporation of structure-level information—particularly quaternary structure and interface localization—could enhance interpretative consistency. Genetic skeletal disorders offer a particularly informative context for such structure-aware variant interpretation because their phenotypes are often highly specific and reproducible at the radiographic level. In many skeletal dysplasias, distinct combinations of bone morphology, growth plate abnormalities, and skeletal patterning defects provide strong phenotypic signatures that can act as functional readouts of disrupted molecular pathways. Consequently, when variants cluster within structurally defined functional regions—such as catalytic sites, RNA-binding motifs [40], or protein–protein interfaces—the corresponding skeletal phenotype can provide independent support for their functional relevance. In this sense, characteristic radiographic phenotypes may serve as a quasi-functional test, strengthening genotype–structure correlations even in the absence of direct biochemical assays. Conversely, when variants of uncertain significance localize outside known functional regions or fail to correlate with the expected structural mechanism, structural mapping can help prioritize candidates for targeted biochemical or cellular assays. This bidirectional relationship between phenotype specificity and structural context makes genetic skeletal disorders particularly well suited for structure-guided interpretation of missense variants.

Structural mapping can directly inform ACMG criteria such as PM1 (location in a critical functional domain or mutational hotspot) by identifying variants that cluster at catalytic sites or obligate protein–protein interfaces. In multimeric complexes, interface-disrupting variants may plausibly exert dominant-negative effects or impair complex assembly, mechanisms that are not readily inferred from sequence conservation alone. In cases where experimental data demonstrate interface sensitivity—such as VPS33A residues required for VPS16 recruitment—structural context may also support PS3 (functional evidence) when mutagenesis studies validate disruption of binding.

Importantly, this approach is particularly valuable in ultra-rare Mendelian disorders, where recurrence-based evidence is often unavailable and cohort-level statistical power is limited. By mapping variants onto experimentally resolved or high-confidence predicted structures, clinicians and molecular geneticists can generate mechanistic hypotheses, prioritize variants of uncertain significance for functional testing, and contextualize rare missense substitutions within three-dimensional architecture. In multimeric assemblies such as the MCM helicase or BBSome complex, interpreting variants across subunits simultaneously may reveal shared catalytic or interface-sensitive regions that would be missed under a monomer-centric framework, potentially enabling interpretation of variants in complex subunits not previously associated with GSDs. In this context, spatial clustering of variants across interacting subunits can reveal structurally equivalent perturbations, where distinct mutations converge on disruption of the same catalytic surface, interaction interface, or regulatory motif. Recognition of such structural equivalence provides a rational framework to transfer pathogenicity inference from known pathogenic variants to nearby VUS within the same structural region.

We do not propose that structural mapping could replace existing classification systems. Rather, our findings support the incorporation of standardized structure-aware annotations into variant interpretation pipelines, ideally as a complementary layer of evidence. As structural coverage continues to expand through experimental efforts and improved predictive models, systematic integration of protein architecture, interface topology, and complex-level organization into clinical genomics workflows may help reduce the burden of variants of uncertain significance and strengthen genotype–phenotype inference in rare disease diagnostics.

Several limitations of this study should be acknowledged. Our analysis relies on existing databases and annotations, which are incomplete and continuously evolving. Both experimental structures and AF2 models represent largely static views of proteins and do not capture the full spectrum of conformational dynamics that often underlie biological function. Multimeric states were inferred from available experimental structures rather than validated across physiological conditions. In addition, the overlap with cancer-associated genes was based on publicly available resources and does not fully capture tiered or context-specific classifications. These limitations highlight areas where future work, including integration of conformational dynamics, experimental validation, and expanded clinical datasets, will be essential.

Structural analyses have traditionally focused either on individual disease genes or on large pan-cancer mutation datasets. In contrast, systematic structural mapping of all variants associated with a specific disease class has rarely been attempted. Our analysis of genetic skeletal disorders demonstrates that such an approach can reveal recurring structural mechanisms shared across diverse molecular systems. Future directions include extending this structure-aware framework to incorporate protein dynamics, systematic interface scoring, and automated integration into clinical variant interpretation pipelines. Expanding the approach to other rare disease groups may help uncover shared molecular principles and facilitate cross-disease learning. Ultimately, bridging genomic data with protein structure and function will be critical for advancing precision medicine in both rare and common diseases.

## Conclusions

In this study, we provide the first comprehensive protein-level landscape of genes associated with genetic skeletal disorders, integrating structural coverage, predicted models, multimeric assemblies, biological function, and clinical variant data. Our results demonstrate that structural knowledge across GSD proteins is uneven and often incomplete, limiting interpretation of genetic variation when considered at the sequence level alone. AlphaFold2 models substantially extend structural coverage but require careful contextualization, particularly for multimeric and interaction-dependent proteins.

Structural analyses have traditionally focused either on individual disease genes or on large pan-cancer mutation datasets. In contrast, systematic structural mapping of all variants associated with a specific disease class has rarely been attempted. Our analysis of genetic skeletal disorders demonstrates that such an approach can reveal recurring structural mechanisms shared across diverse molecular systems.By highlighting the importance of protein complexes, interaction interfaces, and shared structural and molecular mechanisms across disease contexts, this work underscores the value of structure-aware genomics for rare disease research. The framework presented here provides a foundation for improving variant interpretation, prioritizing uncertain variants, and generating mechanistic hypotheses in genetic skeletal disorders. More broadly, it illustrates how integrating protein structure into genomic analyses can enhance our understanding of genotype–phenotype relationships and support the development of more precise and informative diagnostic strategies

## Supplementary Figures

**Supplementary Fig. 1.**
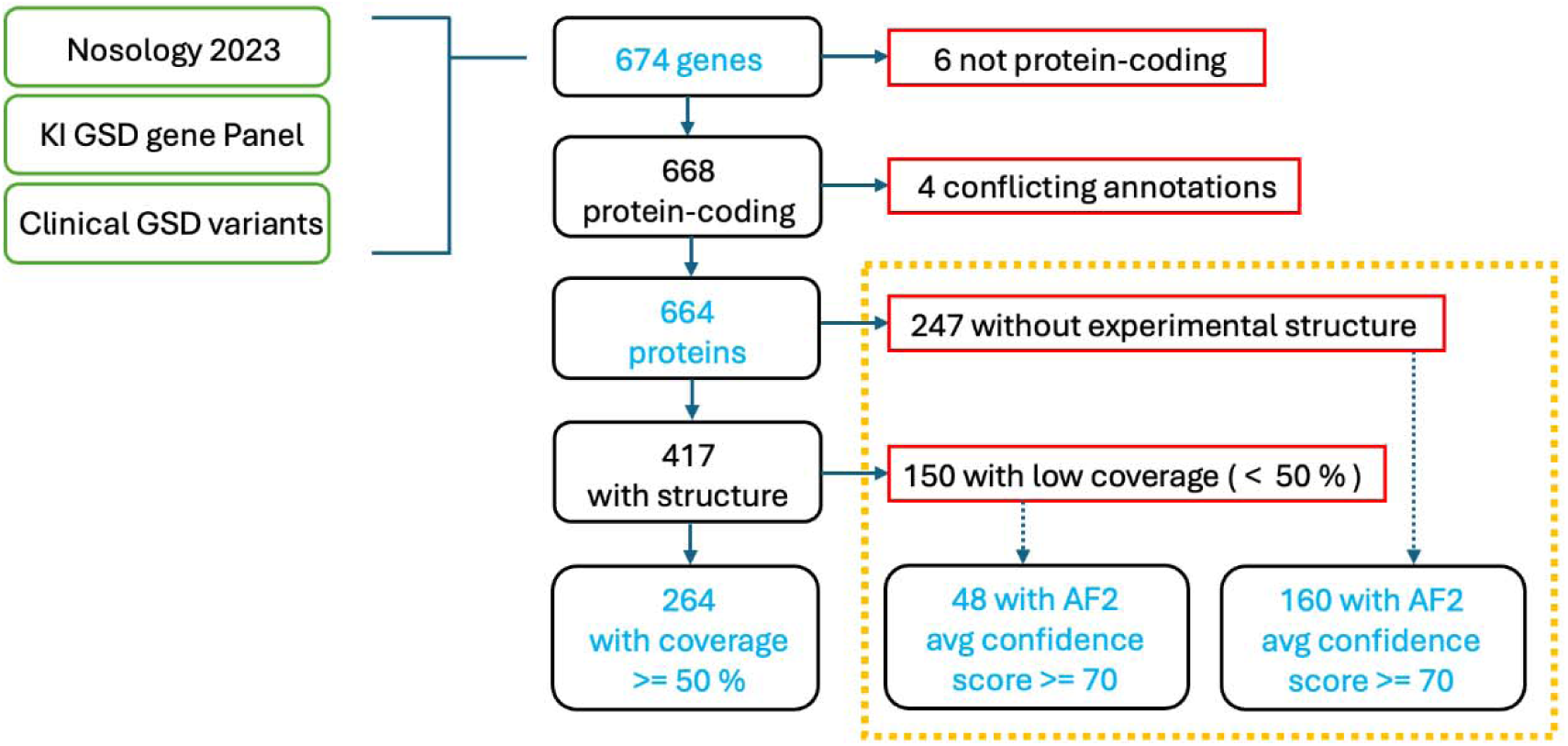
Overview of structural knowledge statistics across GSD-associated proteins. Summary of experimental structure availability, sequence coverage, AlphaFold2 model confidence, and observed multimeric states across the GSD protein dataset. The figure highlights systematic gaps in structural knowledge and illustrates how predicted models and multimeric context contribute complementary information for variant interpretation.

**Supplementary Fig. 2.**
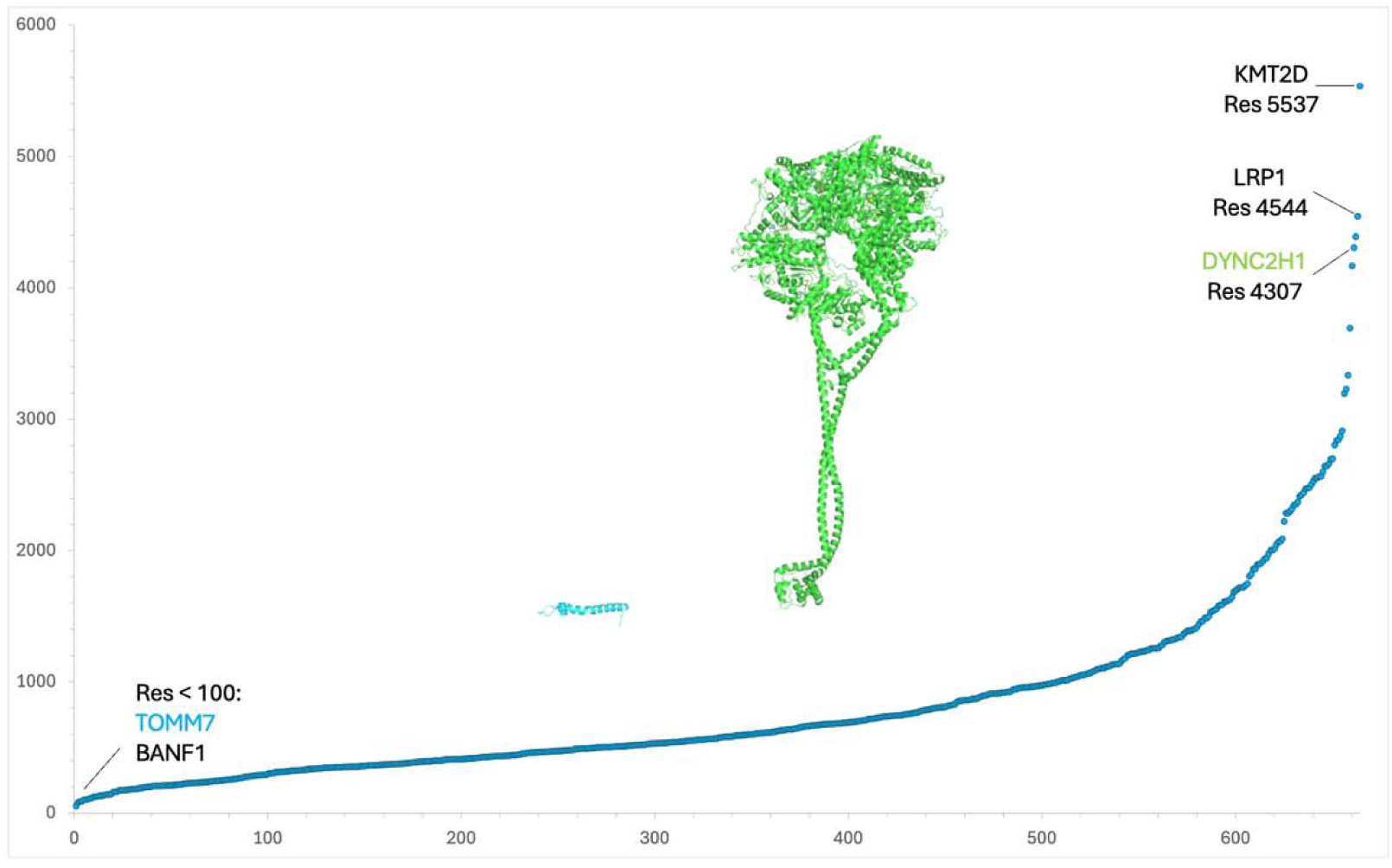
Protein size distribution of genetic skeletal disorder–associated proteins. Distribution of canonical protein lengths for the 664 protein-coding genes included in this study. Protein sizes span more than two orders of magnitude, reflecting the functional and structural heterogeneity of genetic skeletal disorder (GSD) proteins. The inset highlights representative extremes, with DYNC2H1 among the largest proteins and TOMM7 among the smallest, illustrating challenges for experimental structural characterization of large multidomain proteins.

**Supplementary Fig. 3.**
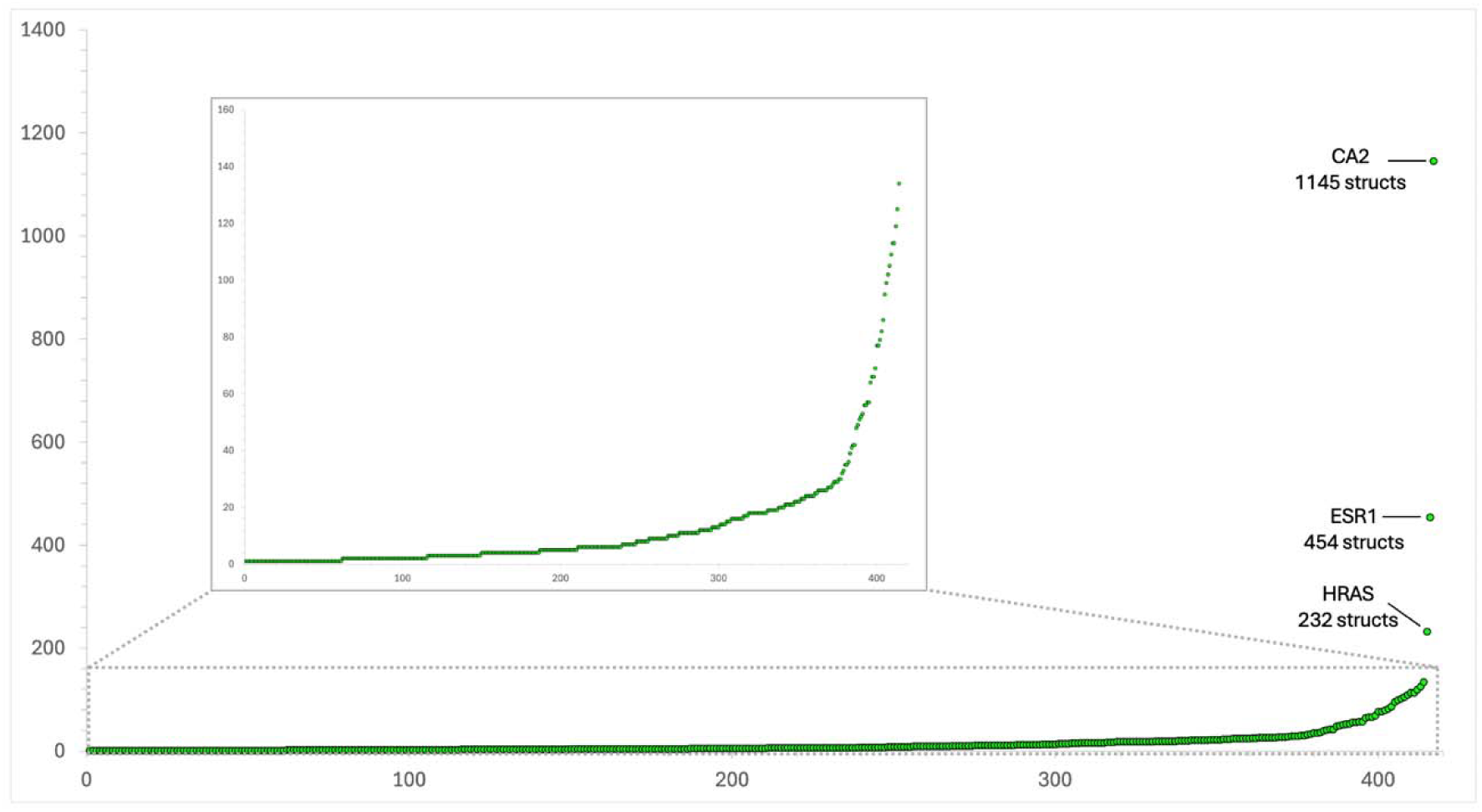
Availability of experimental protein structures across GSD-associated genes. Number of experimentally determined structures per GSD-associated protein based on Protein Data Bank entries. While a subset of proteins shows extensive structural representation—often due to intensive study in non-skeletal disease contexts—the majority of proteins are represented by relatively few structures, and 37% lack experimental structures entirely. The inset shows proteins represented by fewer than 200 structures.

**Supplementary Fig. 4.**
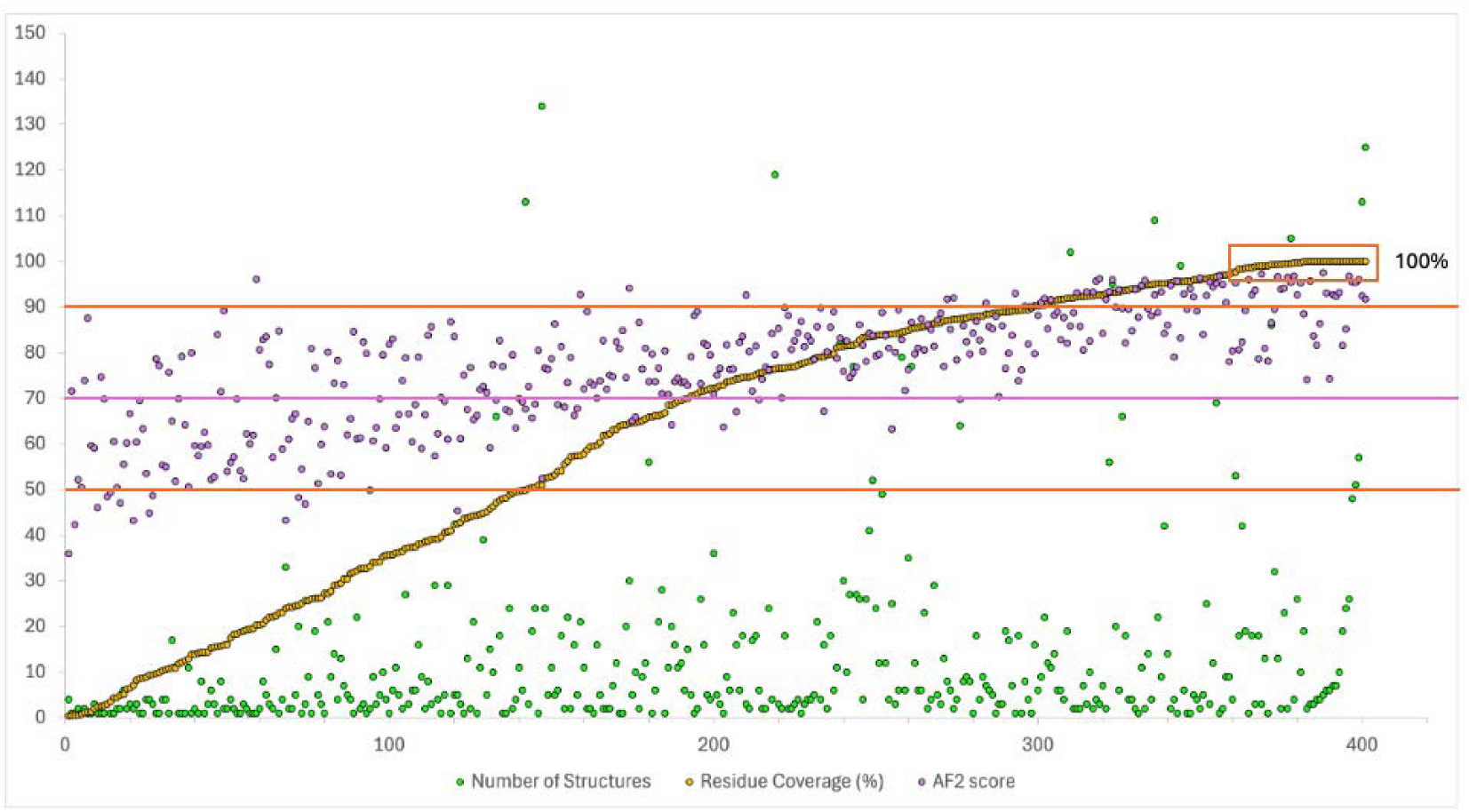
Sequence coverage of GSD proteins by experimental structures. Distribution of experimental sequence coverage, defined as the fraction of residues in the canonical protein sequence observed in at least one experimental structure. Many proteins exhibit partial or fragmentary coverage, limiting structure-based interpretation of genetic variants to specific domains rather than full-length architectures. Overlap between the number of experimental structures and AlphaFold2 average confidence score (AF2 score). Lines are depicted for 50% and 90% for the coverage and 70% for AF2 score. The square includes structures with 100% coverage. Proteins with more than 200 structures are not included.

## Supplementary Tables

Large datasets are provided as separate supplementary files (Additional Files 1–5), while smaller tables are included in the Supplementary Information document.

**Supplementary Table S1.** Gene lists integrated in this study, including the Karolinska University Hospital GSD diagnostic panel (KI_PAN), Nosology 2023 (NOS23), and clinically investigated GSD genes (KI_CLI).

| Datasets | Abbreviation | Description | Reference |
| --- | --- | --- | --- |
| **Karolinska Hospital Gene Panel** | KI_Pan | Genes used in the screening panel in clinical practice at Karolinska Hospital | n.a. |
| **Karolinska Hospital Clinical Cases** | KI_Cli | Pathology-driving genes found in GSD patients at Karolinska Hospital in 2015–2022 | <sup>1</sup> |
| **GSD Nosology 2023** | NOS23 | List of genes included in the later nosology paper for genetic skeletal disorders of 2023 | <sup>2</sup> |

<sup>1</sup> Manuscript in preparation, Lindelöf et al 2026
<sup>2</sup> Unger S et al. *Am J Med Genet A*. 2023;191:1164–209.
n.a. = not applicable

**Supplementary Table S2.**
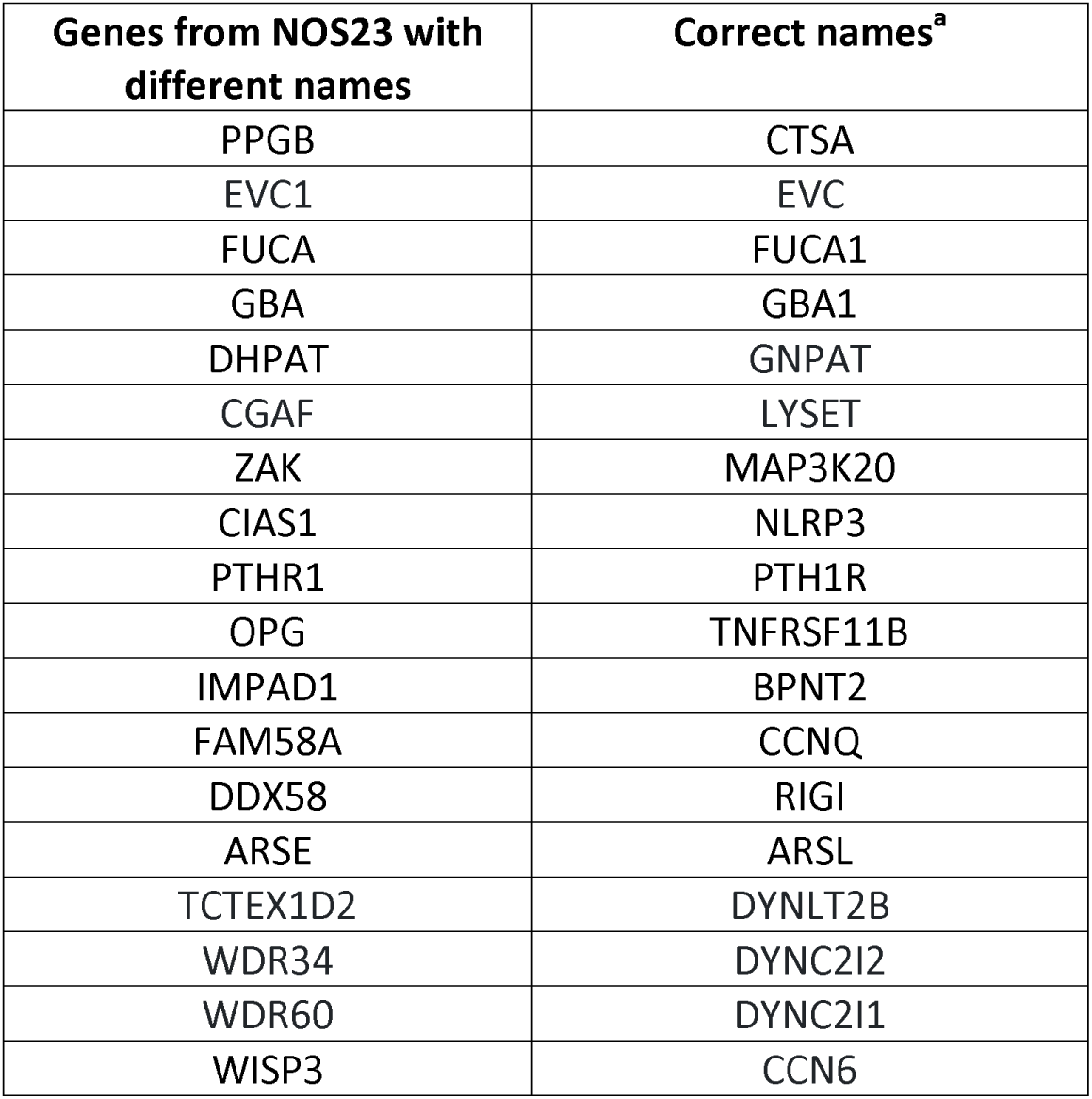
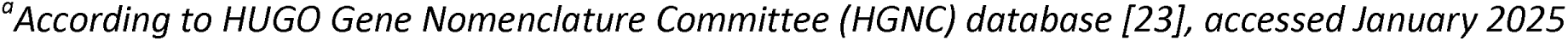
Genes with discrepant nomenclature between Nosology 2023 and official HGNC gene symbols, with manual resolution.

| Genes from NOS23 with different names | Correct names<sup>a</sup> |
| --- | --- |
| PPGB | CTSA |
| EVC1 | EVC |
| FUCA | FUCA1 |
| GBA | GBA1 |
| DHPAT | GNPAT |
| CGAF | LYSET |
| ZAK | MAP3K20 |
| CIAS1 | NLRP3 |
| PTHR1 | PTH1R |
| OPG | TNFRSF11B |
| IMPAD1 | BPNT2 |
| FAM58A | CCNQ |
| DDX58 | RIGI |
| ARSE | ARSL |
| TCTEX1D2 | DYNLT2B |
| WDR34 | DYNC2I2 |
| WDR60 | DYNC2I1 |
| WISP3 | CCN6 |

<sup>a</sup>According to HUGO Gene Nomenclature Committee (HGNC) database [23], accessed January 2025

**Supplementary Table S3.** Excluded NOS23 entries corresponding to loci, regulatory elements, or non–protein-coding annotations.

| Genes or loci | Reason for exclusion |
| --- | --- |
| 10q24 | Locus encompassing several genes |
| 11p15.5 | Locus encompassing several genes |
| 14q32 | Locus encompassing several genes |
| 2p14 - p13.3 | No gene identified |
| 2q35 - 36 | Locus deletion involved in abnormal PAX3 expression (PAX3 is included in the study) |
| 8q22.1 | Locus encompassing several genes |
| MAENLI | lncRNA regulating EN1 (EN1 is included in the study) |
| ZRS | Regulatory element for SHH gene in intron of LMBR1 gene (both SHH and LMBR1 genes are included in the study) |

**Supplementary Table S4.** Additional File 1. Final curated list of GSD-associated genes with HGNC identifiers, UniProt accessions, and primary gene names.

**Supplementary Table S5.** Excluded genes due to non–protein-coding status or unresolved protein–gene mapping conflicts.

| Gene Name | HGNC id | Reason for exclusion |
| --- | --- | --- |
| HAGLR | 43755 | non protein-coding |
| MIR140 | 31527 | non protein-coding |
| MIR17HG | 23564 | non protein-coding |
| RMRP | 10031 | non protein-coding |
| RNU12 | 19380 | non protein-coding |
| RNU4ATAC | 34016 | non protein-coding |
| GNAS | 4392 | Conflicting gene-protein mapping |
| RAB34 | 16519 | Conflicting gene-protein mapping |
| VCP | 12666 | Conflicting annotations unrelated to GSD |

**Supplementary Table S6.** Additional File 1. Final dataset of 664 protein-coding GSD genes included in structural and functional analyses.

**Supplementary Table S7.** Additional File 2. Protein structural statistics, including canonical sequence length, number of experimental structures, sequence coverage, AlphaFold2 model availability, and average pLDDT confidence scores.

**Supplementary Table S8.**
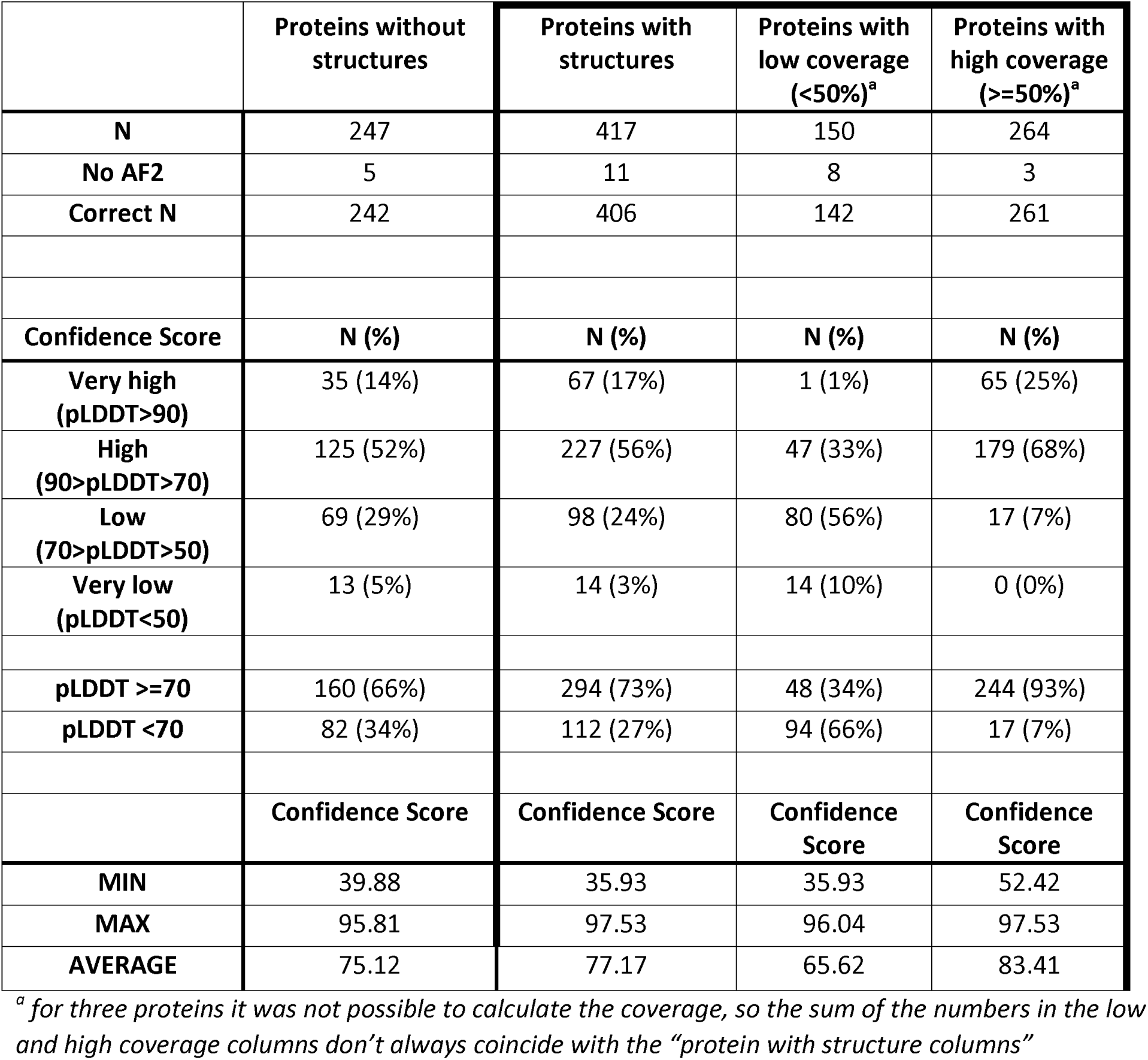
AlphaFold2 confidence scores for GSD proteins with and without experimental structures.

**Supplementary Table S9.**
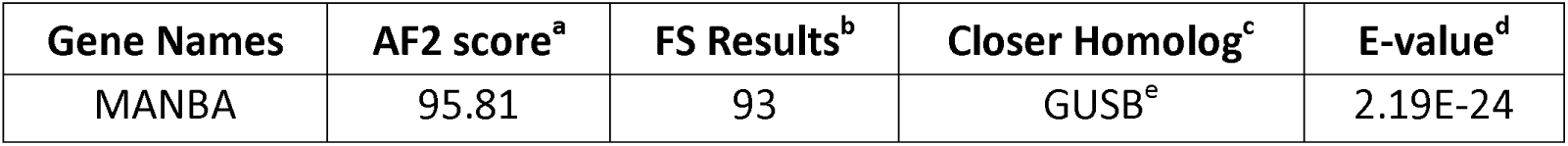
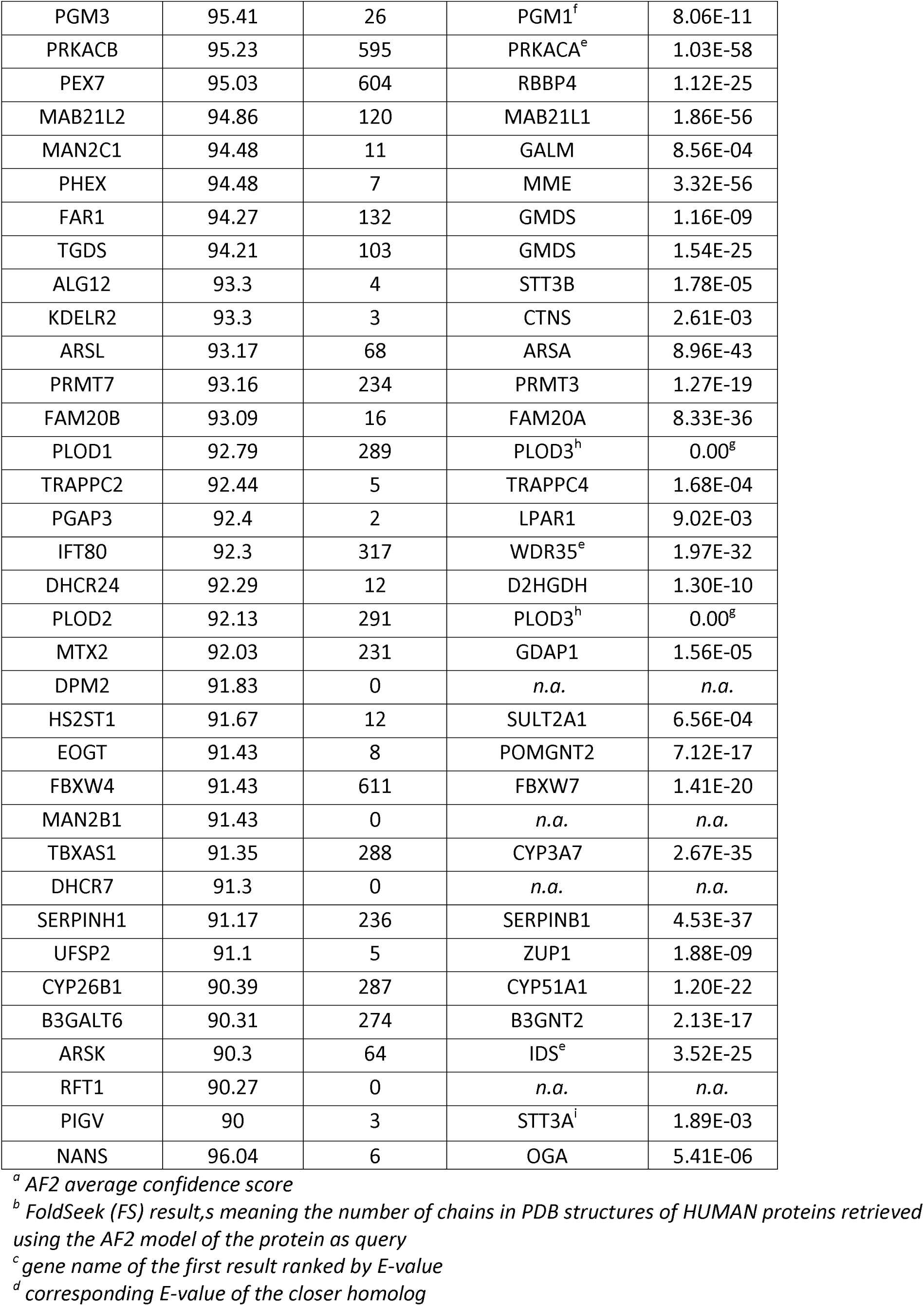
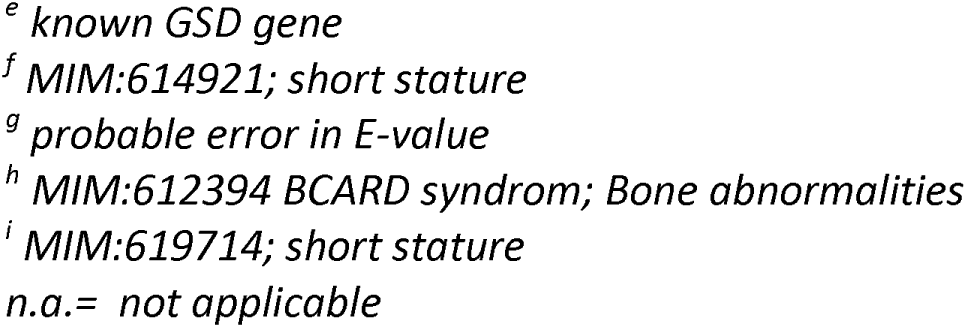
High-confidence AlphaFold2 models for structural homologs of GSD proteins without experimental structures or with low experimental coverage (“rescued” proteins).

**Supplementary Table S10.** Additional File 3. Annotated biological pathways associated with GSD proteins

**Supplementary Table S11.** Additional File 3. GSD Genes associated with statistically significant pathways

**Supplementary Table S12.** Additional File 4. Observed multimeric states of GSD proteins based on experimental structures, including monomeric and oligomeric assemblies.

**Supplementary Table S13.** Additional File 4. Protein co-occurrence analysis showing GSD protein partners captured within the same experimental structures.

**Supplementary Table S14.**
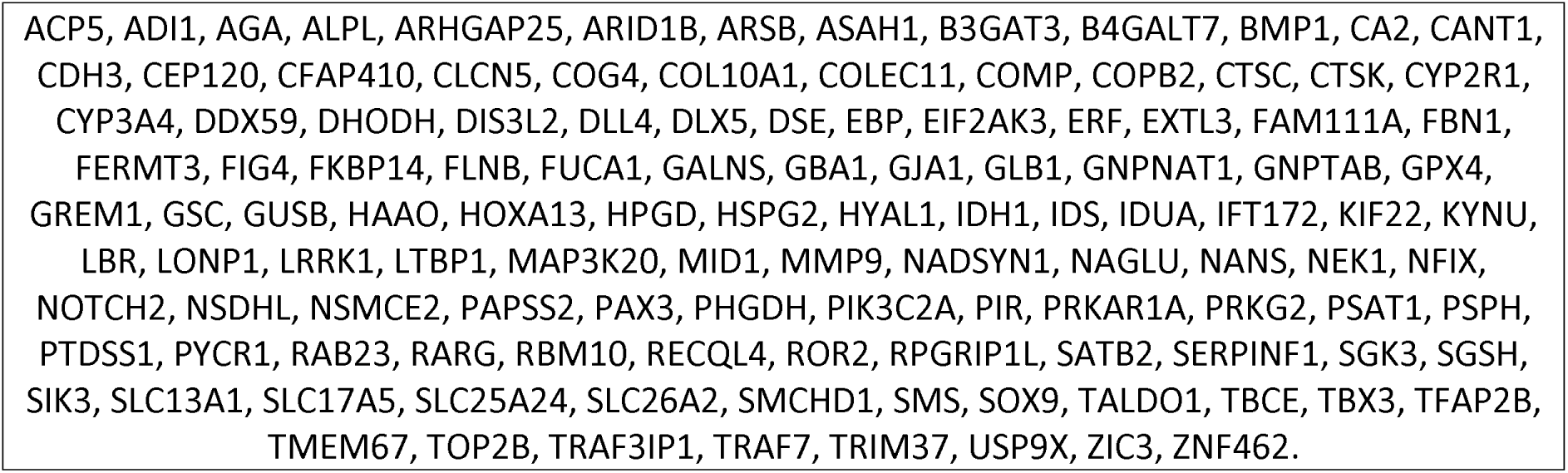
Proteins without partners in the experimental structures.

**Supplementary Table S15.**
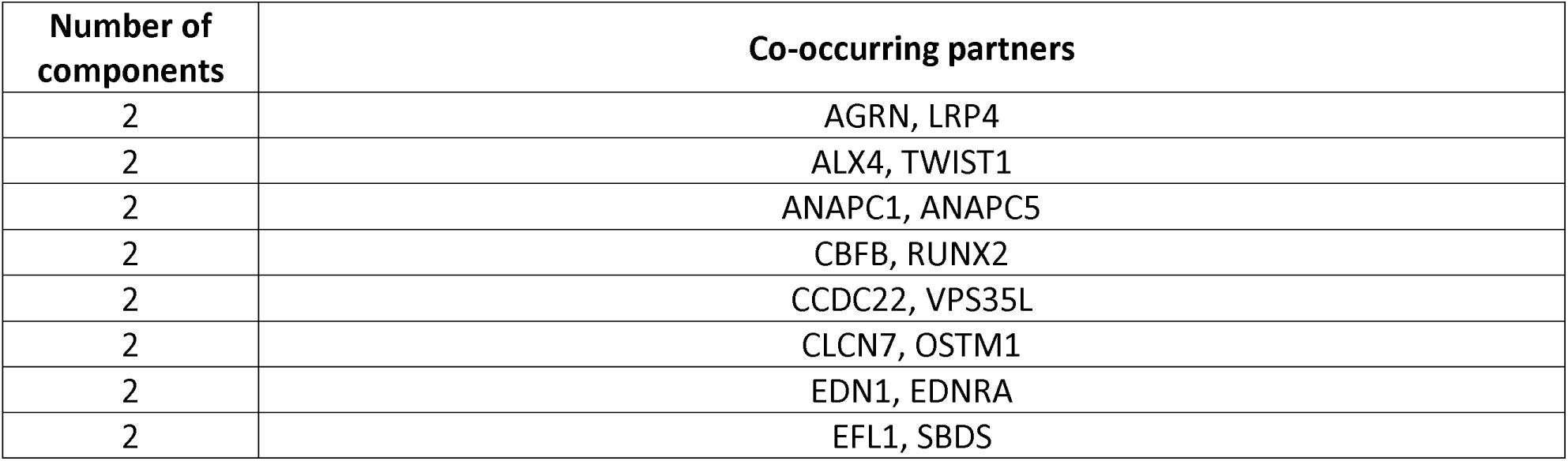
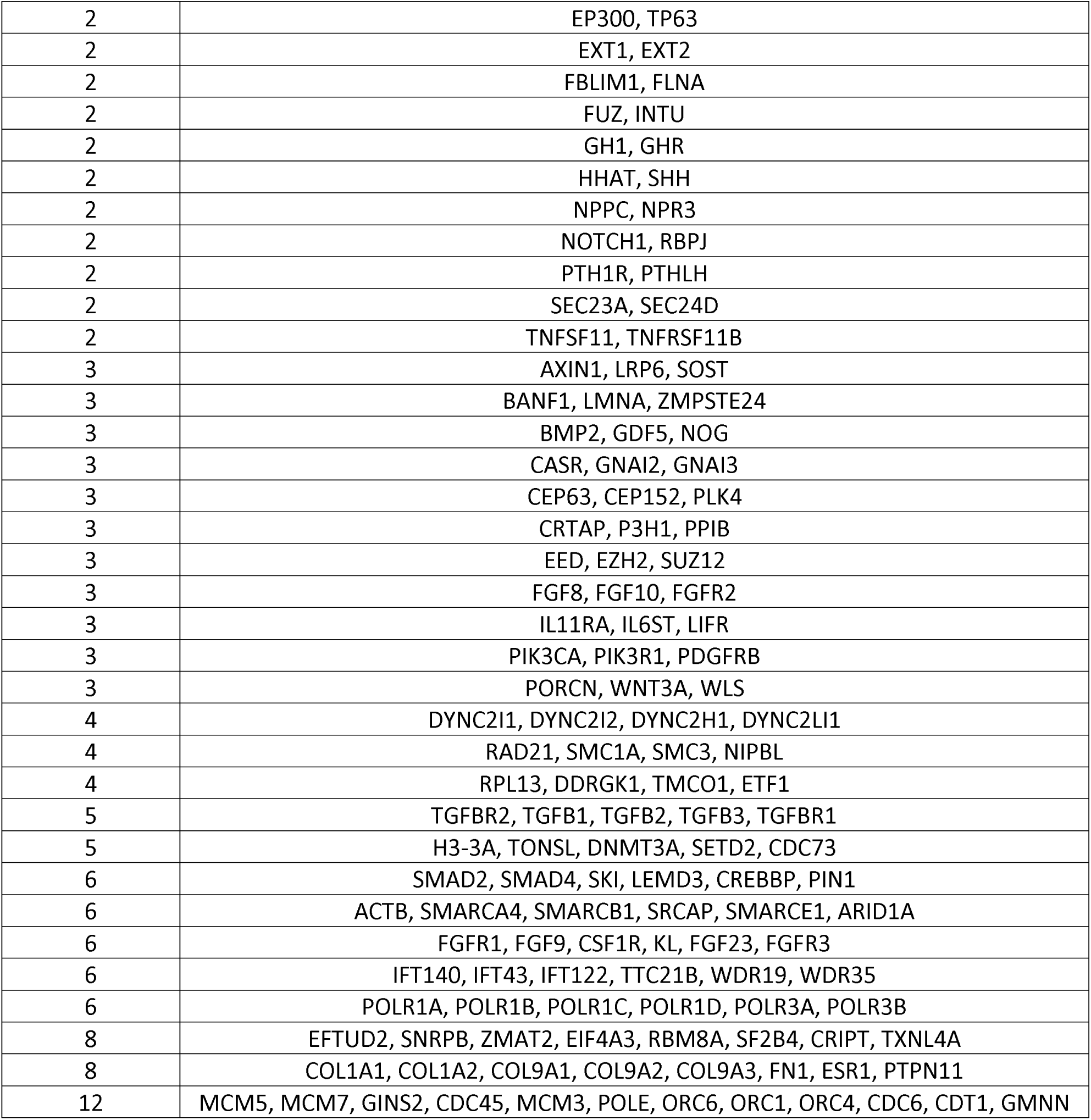
Protein co-occurrence analysis showing GSD–GSD protein partners.

**Supplementary Table S16.**
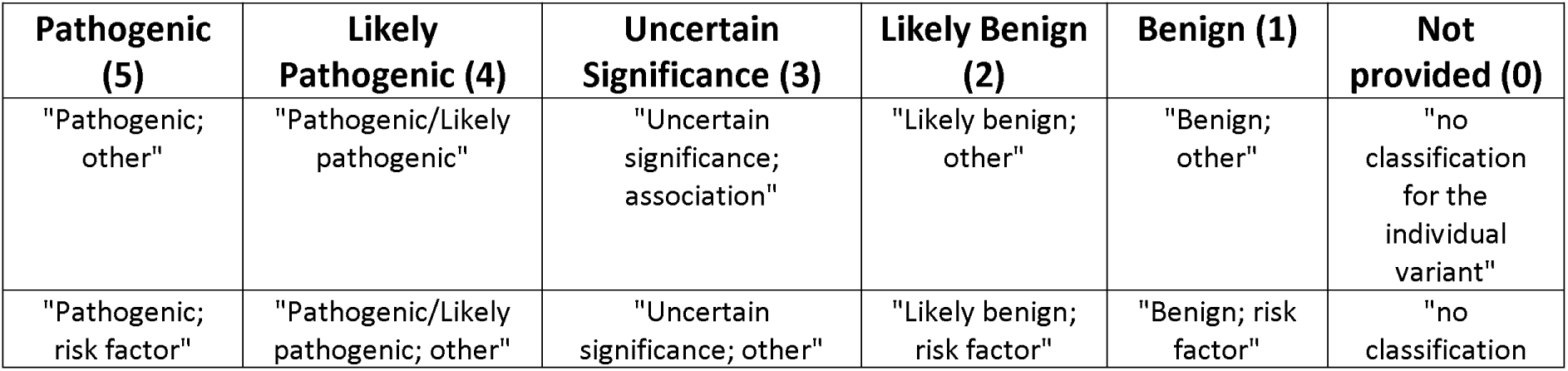
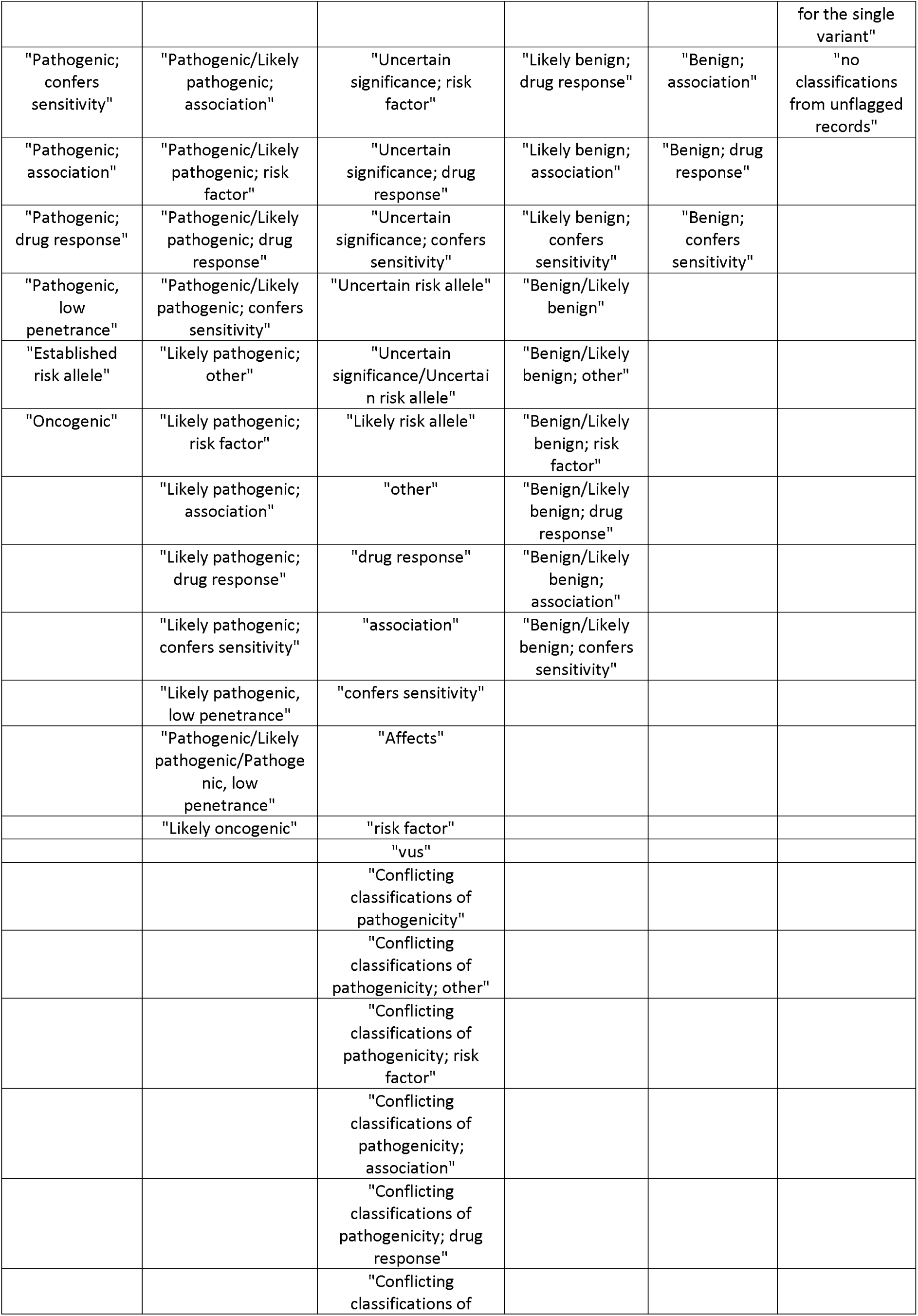
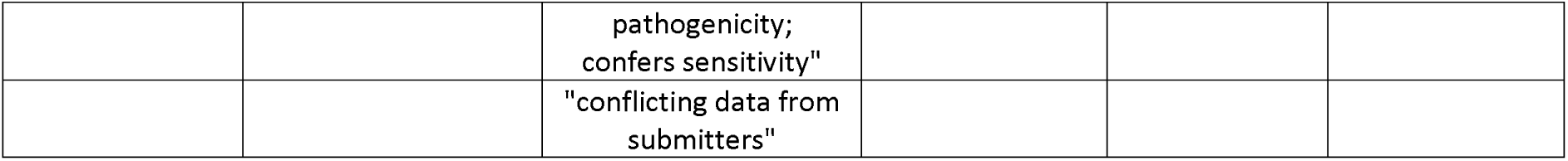
ClinVar pathogenicity re-classification.

**Supplementary Table S17.** Subset of GSD proteins selected for detailed clinical variant mapping, including criteria based on sequence coverage (>= 95 %), protein length (>=400), and structural representation (<= 3).

| Gene Name | Number of Structures | Residue Coverage | Residue Coverage (%) | Partners? | GSD Partners? |
| --- | --- | --- | --- | --- | --- |
| BBS1 | 1 | 567 | 95.6 | YES |  |
| EFL1 | 2 | 1120 | 100 | YES | SBDS |
| HYAL1 | 1 | 416 | 95.6 | NO |  |
| KYNU | 3 | 448 | 96.3 | NO |  |
| NADSYN1 | 1 | 696 | 98.6 | NO |  |
| NAGLU | 1 | 720 | 96.9 | NO |  |
| SGSH | 2 | 482 | 96 | NO |  |
| TTC21B | 3 | 1316 | 100 | YES | IFT140, IFT43,<br>IFT122,<br>WDR19,<br>WDR35 |
| VPS33A | 2 | 592 | 99.3 | YES |  |

**Supplementary Table S18.** Additional File 5. Overlap between GSD-associated genes and cancer-associated genes from publicly available Cancer Gene Census resources.

## Supplementary Information

**Additional file 1** Tables S4 & S6. Gene curation and protein dataset definition.

*Supplementary Table S4. Final curated list of GSD-associated genes with HGNC identifiers, UniProt accessions, and primary gene names*.

*Supplementary Table S6. Final dataset of 664 protein-coding GSD genes included in structural and functional analyses*.

**Additional file 2** Table S7. Structural statistics and AlphaFold2 confidence scores.

*Supplementary Table S7. Structural statistics for GSD proteins, including canonical sequence length, number of experimental structures, sequence coverage, AlphaFold2 model availability, and average pLDDT confidence scores*.

**Additional file 3** Tables S10–S11. Pathway enrichment analysis of GSD proteins.

Supplementary Table S10. Annotated biological pathways associated with GSD proteins identified using PANTHER pathway analysis.

Supplementary Table S11. GSD genes associated with statistically significant pathways.

**Additional file 4** Tables S12–S13. Multimeric assemblies and protein co-occurrence.

Supplementary Table S12. Observed multimeric states of GSD proteins based on experimental structures.

Supplementary Table S13. Protein co-occurrence analysis showing interacting partners captured within the same experimental structures.

**Additional file 5** Table S18. Overlap with cancer-associated genes.

*Supplementary Table S18. Overlap between GSD-associated genes and cancer-associated genes derived from the Cancer Gene Census dataset*.

## List of Abbreviations

ACMG: American College of Medical Genetics and Genomics
AMP: Association for Molecular Pathology
AF2: AlphaFold2
AF3: AlphaFold3
AFDB: AlphaFold Protein Structure Database
CGC: Cancer Gene Census
ECM: Extracellular Matrix
FGF: Fibroblast Growth Factor
GNAS: Guanine Nucleotide-Binding Protein Alpha Subunit
GO: Gene Ontology
GPCR: G Protein-Coupled Receptor
GSD: Genetic Skeletal Disorders
HGNC: HUGO Gene Nomenclature Committee
HOPS: Homotypic Fusion and Protein Sorting Complex
KI_CLI: Karolinska University Hospital Clinical GSD Gene Set
KI_PAN: Karolinska University Hospital GSD Diagnostic Gene Panel
LOF: Loss of Function
MANE: Matched Annotation from NCBI and EMBL-EBI
MCM: Minichromosome Maintenance (Helicase) Complex
MD: Molecular Dynamics
NOS23: Nosology of Genetic Skeletal Disorders 2023 Revision
PDB: Protein Data Bank
pLDDT: Predicted Local Distance Difference Test
RTK: Receptor Tyrosine Kinase
SIFTS: Structure Integration with Function, Taxonomy and Sequences
SMAD: Mothers Against Decapentaplegic (TGF-β signaling mediators)
TF: Transcription Factor
UniProt: Universal Protein Resource
VUS: Variant of Uncertain Significance
WGS: Whole Genome Sequencing

## Declarations

### Ethics approval and consent to participate

Not Applicable

### Consent for publication

Not Applicable

### Availability of data and materials

The original data are available in the databases used (ClinVar, PDB).

Processed structural coverage statistics, AlphaFold confidence metrics, multimeric annotations, and cancer-gene overlap data are provided in the Supplementary Information. Additional processed datasets and code are available from the corresponding author upon reasonable request.

## Authors’ contributions

GG and LO conceptualized the study, acquired the funding, interpreted the data and wrote the manuscript. SGP developed the tools, generated, analyzed and interpreted the data and wrote the manuscript. All authors have read, edited, and approved the final manuscript.

## Acknowledgements

Thanks to Anna Hammarsjö for the list of genes from KI GSD panel, to Hillevi Lindelöf for the list of KI clinical variants and to Domenico Scaramozzino, Byung Ho Lee and Michael Nagy for the helpful advice.

## References

1. Geister KA, Camper SA. Advances in Skeletal Dysplasia Genetics. Annu Rev Genomics Hum Genet. 2015;16:199–227. 10.1146/annurev-genom-090314-045904

2. Grigelioniene G, Nishimura G. Exploring human genetic skeletal disorders provides important insights into skeletogenesis and elucidates basic developmental signaling pathways. EBioMedicine. 2020;62. 10.1016/j.ebiom.2020.103091

3. Smith CIE, Bergman P, Hagey DW. Estimating the number of diseases – the concept of rare, ultra-rare, and hyper-rare. iScience [Internet]. Elsevier; 2022 [cited 2022 Sep 1];25:104698. 10.1016/J.ISCI.2022.104698

4. Stenson PD, Mort M, Ball EV, Chapman M, Evans K, Azevedo L, et al. The Human Gene Mutation Database (HGMD®): optimizing its use in a clinical diagnostic or research setting. Human Genetics. 2020;139. 10.1007/s00439-020-02199-3

5. Landrum MJ, Lee JM, Benson M, Brown GR, Chao C, Chitipiralla S, et al. ClinVar: Improving access to variant interpretations and supporting evidence. Nucleic Acids Research. 2018;46. 10.1093/nar/gkx1153

6. Ng PC, Henikoff S. SIFT: Predicting amino acid changes that affect protein function. Nucleic Acids Res. 2003;31(13):3812–3814. doi:10.1093/nar/gkg509

7. Schwarz JM, Rödelsperger C, Schuelke M, Seelow D. MutationTaster evaluates disease-causing potential of sequence alterations. Nature Methods. 2010;7. 10.1038/nmeth0810-575

8. Ferrer-Costa C, Gelpí JL, Zamakola L, Parraga I, de la Cruz X, Orozco M. PMUT: A web-based tool for the annotation of pathological mutations on proteins. Bioinformatics. 2005;21. 10.1093/bioinformatics/bti486

9. Rogers MF, Shihab HA, Mort M, Cooper DN, Gaunt TR, Campbell C. FATHMM-XF: Accurate prediction of pathogenic point mutations via extended features. Bioinformatics. 2018;34. 10.1093/bioinformatics/btx536

10. Orellana L. Are Protein Shape-Encoded Lowest-Frequency Motions a Key Phenotype Selected by Evolution? Applied Sciences [Internet]. Multidisciplinary Digital Publishing Institute; 2023 [cited 2023 Sep 3];13:6756. 10.3390/app13116756

11. Orellana L. Large-Scale Conformational Changes and Protein Function: Breaking the in silico Barrier. Frontiers in Molecular Biosciences [Internet]. 2019 [cited 2023 Oct 13];6. https://www.frontiersin.org/articles/10.3389/fmolb.2019.00117. Accessed 13 Oct 2023

12. Khanna T, Hanna G, Sternberg MJE, David A. Missense3D-DB web catalogue: an atom-based analysis and repository of 4M human protein-coding genetic variants. Human Genetics. 2021;140. 10.1007/s00439-020-02246-z

13. Ofoegbu TC, David A, Kelley LA, Mezulis S, Islam SA, Mersmann SF, et al. PhyreRisk: A Dynamic Web Application to Bridge Genomics, Proteomics and 3D Structural Data to Guide Interpretation of Human Genetic Variants. Journal of Molecular Biology. 2019;431. 10.1016/j.jmb.2019.04.043

14. Laskowski RA, Stephenson JD, Sillitoe I, Orengo CA, Thornton JM. VarSite: Disease variants and protein structure. Protein Science. 2020;29. 10.1002/pro.3746

15. Cheng J, Novati G, Pan J, Bycroft C, Žemgulytė A, Applebaum T, et al. Accurate proteome-wide missense variant effect prediction with AlphaMissense. Science [Internet]. American Association for the Advancement of Science; 2023 [cited 2024 Feb 27];381:eadg7492. 10.1126/science.adg7492

16. Mhashal AR, Yoluk O, Orellana L. Exploring the conformational impact of novel glycine receptor mutations through coarse-grained analysis and atomistic simulations. Frontiers in Molecular Biosciences. 2022;9:890851.

17. Abramson J, Adler J, Dunger J, Evans R, Green T, Pritzel A, et al. Accurate structure prediction of biomolecular interactions with AlphaFold 3. Nature. 2024;630:493–500. 10.1038/s41586-024-07487-w

18. Jumper J, Evans R, Pritzel A, Green T, Figurnov M, Ronneberger O, et al. Highly accurate protein structure prediction with AlphaFold. Nature [Internet]. Springer US; 2021;596:583–9. 10.1038/s41586-021-03819-2

19. Varga MJ, Richardson ME, Chamberlin A. Structural biology in variant interpretation: Perspectives and practices from two studies. Am J Hum Genet [Internet]. 2025 [cited 2025 Dec 16];112:984–92. 10.1016/j.ajhg.2025.03.010

20. Yang Z, Zeng X, Zhao Y, Chen R. AlphaFold2 and its applications in the fields of biology and medicine. Sig Transduct Target Ther [Internet]. Nature Publishing Group; 2023 [cited 2025 Dec 16];8:115. 10.1038/s41392-023-01381-z

21. Berman HM, Westbrook J, Feng Z, Gilliland G, Bhat TN, Weissig H, et al. The Protein Data Bank. Allen FH, Berghoff G, Sievers R, editors. Nucleic Acids Research. Research Collaboratory for Structural Bioinformatics (RCSB), Rutgers University, Piscataway, NJ 08854-8087, USA.; 2000;28:235–42.

22. Berman H, Henrick K, Nakamura H. Announcing the worldwide Protein Data Bank. Nat Struct Mol Biol [Internet]. Nature Publishing Group; 2003 [cited 2025 Dec 16];10:980–980. 10.1038/nsb1203-980

23. Unger S, Ferreira CR, Mortier GR, et al. Nosology of genetic skeletal disorders: 2023 revision. Am J Med Genet A. 2023;191(5):1164–1209. doi:10.1002/ajmg.a.63132

24. HUGO Gene Nomenclature Committee at the University of Cambridge (www.genenames.org)

25. UniProt C. UniProt: the Universal Protein Knowledgebase in 2023. Nucleic Acids Res. 2023;51(D1):D523–D31.

26. Ahmad S, da Costa Gonzales L J, Bowler-Barnett E H, Rice D L, Kim M, Wijerathne S, Luciani A, Kandasaamy S, Luo J, Watkins X, Turner E, Martin M J, UniProt Consortium The UniProt website API: facilitating programmatic access to protein knowledge Nucleic Acids Research, gkaf394 (2025)

27. Morales, J., Pujar, S., Loveland, J.E. et al. A joint NCBI and EMBL-EBI transcript set for clinical genomics and research. Nature. 2022 Apr;604(7905):310–315.

28. Varadi M, Anyango S, Deshpande M, Nair S, Natassia C, Yordanova G, et al. AlphaFold Protein Structure Database: massively expanding the structural coverage of protein-sequence space with high-accuracy models. Nucleic Acids Res. 2022;50(D1):D439–D44.

29. Velankar S, Dana JM, Jacobsen J, et al. SIFTS: Structure Integration with Function, Taxonomy and Sequences resource. Nucleic Acids Res. 2013;41(Database issue):D483-D489. doi:10.1093/nar/gks1258

30. Dana JM, Gutmanas A, Tyagi N, et al. SIFTS: updated Structure Integration with Function, Taxonomy and Sequences resource allows 40-fold increase in coverage of structure-based annotations for proteins. Nucleic Acids Res. 2019;47(D1):D482-D489. doi:10.1093/nar/gky1114

31. Cock PA, Antao T, Chang JT, Chapman BA, Cox CJ, Dalke A, Friedberg I, Hamelryck T, Kauff F, Wilczynski B and de Hoon MJL (2009) Biopython: freely available Python tools for computational molecular biology and bioinformatics. Bioinformatics, 25, 1422–1423

